# Oncogenic MAPK pathway activation disrupts Schwann cell fate commitment, inducing congenital and progressive neuropathy in mice

**DOI:** 10.1101/2024.04.24.590951

**Authors:** Elise Marechal, Daniel Aldea, Patrice Quintana, Grégoire Mondielli, Nathalie Bernard-Marissal, Mathias Moreno, Claire El Yazidi, Natacha Broucqsault, Valérie Delague, Lauren A. Weiss, Anne Barlier, Heather C. Etchevers

## Abstract

RASopathies, rare congenital syndromes affecting multiple organ systems, often include peripheral neuropathy of unknown origin. While RASopathy-associated gene variants are proto-oncogenic, the impact of timing and mosaicism on pathogenicity remains poorly understood. Here, we investigate the links between Braf, a key mitogen-activated protein kinase (MAPK) effector, and peripheral neuropathy. By targeting *Braf* p.V600E, an oncogenic variant found in mosaic RASopathies, to embryonic Mpz-expressing cells in mice, we induced a congenital Charcot-Marie-Tooth-like degenerative neuropathy. This phenotype was characterized by hyperplastic nerves, hindlimb weakness, and unexpectedly reduced body size. Constitutively active Braf expanded a Jun+ Schwann cell repair state, impairing myelination and nerve homeostasis. To examine relevance to RASopathies, we differentiated patient-derived stem cells bearing the cardio-facio-cutaneous syndrome-associated *BRAF* p.Q257R variant into Schwann cells. Compared to wild-type controls, CFC-derived cells failed to acquire mature phenotypes, instead exhibiting progenitor or repair-type transcriptional profiles. Our findings implicate somatic mosaicism in the unresolved genetic heterogeneity of neuropathies and expand the candidate gene list for peripheral nerve disorders. Moreover, they reveal a MAPK-dependent mechanism linking neural crest-derived Schwann cell dysfunction to both body growth and nerve homeostasis, providing new insights into the mechanisms in RASopathy-associated neuropathy and potential therapeutic targets.

## Introduction

Many cancers and congenital diseases co-opt and deploy the molecular mechanisms of normal organogenesis in inappropriate contexts. Activation of the mitogen-activated protein kinase (MAPK) intracellular signaling pathway drives dozens of cancers in somatic tissues ^1^. In contrast, RASopathies are heterogeneous developmental syndromes that preferentially affect epidermis and pigmentation; cardiovascular, craniofacial and skeletal systems; and neurological or endocrine functions impacting body size and fertility ^2–5^. Pathogenesis results from variants in at least 24 genes that sustain MAPK signaling to, and nuclear translocation of, the extracellular signal-regulated kinases (ERK1/2) ^6^. Activating variants may be expressed constitutionally or as mosaic lineage-specific (somatic) RASopathies ^7^. Most constitutional and somatic RASopathies associate with specific cancer predispositions, particularly when syndromic ^8,9^.

A limited number of MAPK pathway variants engender this array of overlapping clinical outcomes. Oncogenic variants in *BRAF*, particularly encoding changes to valine (p.V)600, drive aggressive forms of melanoma as well as many other cancers ^1,10^. BRAF p.V600E has never been observed in germline disease because of early embryonic lethality, modeled in mice ^11^. BRAF p.V600E yields an active, monomeric kinase, decoupled from RAS-mediated transduction ^12,13^. In contrast, less-activating *BRAF* variants such as p.Q257R underlie a growing list of congenital RASopathies ^8,14^. These include the germline disorders cardio-facio-cutaneous syndrome (CFC; ORPHA:1340) and Noonan syndrome with (ORPHA:500) or without (ORPHA:648) multiple lentigines. Their common clinical features include reduced postnatal growth, dysmorphic facial features, melanocytic nevi or other pigmented lesions, congenital heart defects, and tumor predispositions ^15–17^.

The mechanisms and triggers of neuropathies in the context of RASopathies remain unexplored. However, recent reports describe CFC and Noonan syndrome patients with early-onset neuropathies characterized by pain, sensory loss and/or muscle wasting ^18–20^, resembling severe, early-onset Charcot-Marie-Tooth (CMT) syndromes ^21,22^. Peripheral nervous system (PNS) function relies entirely on Schwann cells (SC), which provide physical and trophic nerve support while insulating large, fast-conduction axons ^23–25^. The PNS — including SC, perineurial stroma and sensory/autonomic neurons — derives from multipotent embryonic neural crest (NC) cells, which also generate craniofacial and cardiovascular connective tissues, neuroendocrine cells and melanocytes ^25–28^. Mek1, the primary MAP2K activating Erk1/2, is essential for nerve myelination during maturation, but its sustained overactivation in adult SC induces murine neuropathy ^29–32^. Despite broad CMT genetic heterogeneity, with dozens of identified genes, no MAPK effectors have been implicated; however, approximately 20% of CMT patients of all subtypes lack a molecular diagnosis ^33^. Given the shared NC origin of affected cell types in RASopathies, we hypothesized that multipotent NC-derived lineages may be vulnerable to prenatal MAPK hyperactivation, compromising their ability to instruct fate or regulate tissue homeostasis over time.

The goal of this study was to assess systemic and cell-autonomous effects of constitutively active BRAF signaling on NC-derived progenitor differentiation. We found that conditional expression of *Braf* p.V600E in mouse cells expressing the SC gene *Mpz* ^34^ (encoding a homophilic adhesion molecule, myelin protein zero [P0]) led to a peripheral neuropathy in association with induction and maintenance of an ineffective repair SC phenotype. Although nerve myelination initiated after birth as usual, mutant peripheral glia rapidly misregulated hundreds of genes associated with mature SC function, including myelination, and nerves underwent active myelin destruction. Unexpectedly, although endocrine tissues had not been intentionally targeted in this model, mice also underwent developmental delay and remained sexually ambiguous or immature. Furthermore, human induced pluripotent stem cells derived from two teenaged CFC syndrome donors, carrying the most frequent *BRAF* missense variant, p.Q257R, were also incapable of terminal SC differentiation *in vitro*, in contrast to cells from wildtype donors, after upregulating repair SC-specific transcripts. These results suggest that molecularly undiagnosed cases of RASopathies, or endophenotypes such as CMT type 1 or idiopathic cryptorchidism ^35^, may be imputable to somatic mosaicism for oncogenic MAPK pathway gene variants in the NC lineage.

## Results

### Sustained Schwann cell MAPK signaling in causes juvenile locomotor and growth deficits

In order to model constitutive Braf-mediated signaling in progenitors of Schwann cells *in vivo*, we used *Mpz-Cre^+/°^* mice, which transcribe *Mpz* (encoding myelin protein zero, or P0) as early as embryonic day 12 in neural crest cells ^36,37^. Crossed to a floxed *Braf* hotspot V600E allele (*Braf*^V600E^) (**Supplemental Figure S1a**), resulting Mpz*-Cre^+/°^*;*Braf*^V600E/+^ (Mpz-*Braf*) mice were born at expected Mendelian ratios, remaining asymptomatic for the first week. From postnatal day (P)7, all Mpz-*Braf* mutants developed signs of neuropathy including tremors, abnormal hindlimb postural reflexes and trailing legs (**Fig 1a**). They nonetheless showed age-appropriate activity in exploration, feeding and grooming. Correlating with phenotypic onset, mutant sciatic nerves exhibited significantly more Erk1/2 phosphorylation than controls, increasing from early (P5; **Fig 1c, e**) to later (P21; **Fig 1d, e**) stages of SC-mediated myelination, demonstrating congenital and persistent MAPK activation in peripheral glia.

**Figure 1.**
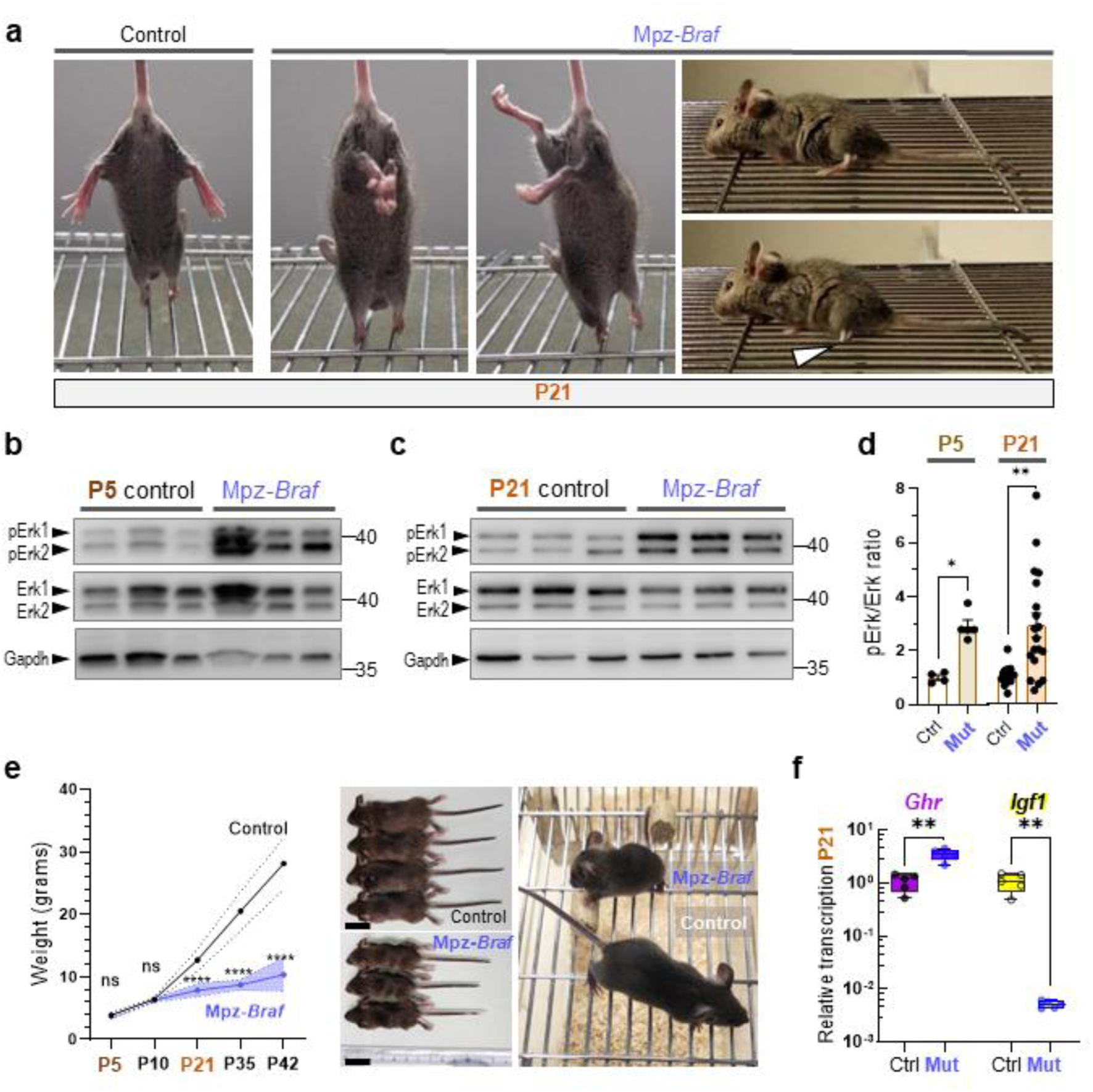
Constitutive MAPK activation mediated by a single copy of *Braf*^V600E^ in murine Schwann cell progenitors induces early peripheral neuropathy and small body size. **a)** By postnatal day (P)21, Mpz-*Braf* mice displayed abnormal hindlimb postural reflexes and ability to keep hindfeet on the bars of a wire cage lid (arrowheads), indicating a functional peripheral nerve defect. **b, c)** Western blot analysis of total and phosphorylated (p)Erk1/2 signaling performed on sciatic nerve lysates from **(b)** P5 and **(c)** P21 control and Mpz-*Braf* mice. Gapdh was detected as a loading control. **d)** Protein densitometry in control (n=4) *vs.* mutant (n=5) sciatic nerve lysates at P5, and control (n=13) *vs.* mutant (n=15) lysates at P21. **e**) Body weight of both sexes was monitored at P5, P10, P21, P35 and P42 in control and Mpz-*Braf* mice, respectively (n = 28 *vs.* 8 at P5, n = 15 *vs.* 5 at P10, n = 54 *vs.* 45 at P21, n = 14 *vs.* 10 at P35, n = 5 *vs.* 5 at P42). **f**) Quantitative PCR analysis of *Ghr* and *Igf1* expression in control (n=5) *vs.* mutant (n=5) livers at P21. Statistical significance from unpaired Mann-Whitney test in **d**), two-way ANOVA followed by Šidák’s post-hoc test in **f**), and Student’s t-test in **g**): ns, not significant; * *p* < 0.05; ** *p* < 0.01; *** *p* < 0.001; **** *p* < 0.0001.

After P10, mutant mice were also visibly smaller than control littermates, with significantly reduced but proportional nose-to-tail length and weight (**Fig 1f**). Since activating *BRAF* mutations have been described in the context of human hypopituitarism ^2^, we looked for a possible central deficiency in growth hormone (Gh) by comparing the transcriptomes of pituitaries at P5 between control and Mpz-*Braf* mice with bulk RNA-sequencing (RNAseq). Gh-releasing hormone receptor *Ghrhr* or *Gh* transcripts, along with all other known coding genes, were not significantly altered in mutants at this stage (**Supplemental Figure S1b**; **Supplemental Table S1**). Gh binds to peripheral Ghr, a type I cytokine receptor, to regulate production of its effector, insulin-like growth factor 1 (Igf1) in the liver and other sites, exerting thereby anabolic effects on bone and muscle ^38^. We therefore compared the transcription of *Ghr* and *Igf1* by RT-qPCR in the livers of P21 Mpz-*Braf* and control littermate mice. While *Ghr* was upregulated on average by 3.3-fold in mutant livers, Igf1 was far more significantly diminished, by 211-fold (**Fig 1g**). The juvenile Mpz-*Braf* growth deficiency was the likely consequence of a near-complete loss of peripheral Igf1 production.

### *Braf^V600E/+^* induces widespread peripheral nerve demyelination and axonal loss

Strikingly broad nerves were visible at autopsy in all Mpz-*Braf* mice at P5 or P21, lacking the structural bands of Fontana ^39^ (**Fig 2a**; **Supplemental Figure S1**). Despite their smaller body size, mutant sciatic nerves were significantly wider at P10 (0.48 ± 0.07 *vs.* 0.31 ± 0.05 mm diameter in controls; **Fig 2b**). This discrepancy stabilized after weaning, with mutant nerves measuring 0.74 ± 0.09 mm at P21 (*vs.* 0.39 ± 0.05 mm in controls), and 0.78 ± 0.05 mm at P35 (*vs.* 0.50 ± 0.04 mm). Nerve enlargement was consistent across all examined peripheral nerves in P21 Mpz-*Braf* mice, including sensorimotor (peroneal, tibial, sural, trigeminal, ophthalmic) and sensory (saphenous, maxillary) nerves (**Supplemental Figure S1c-e**).

**Figure 2.**
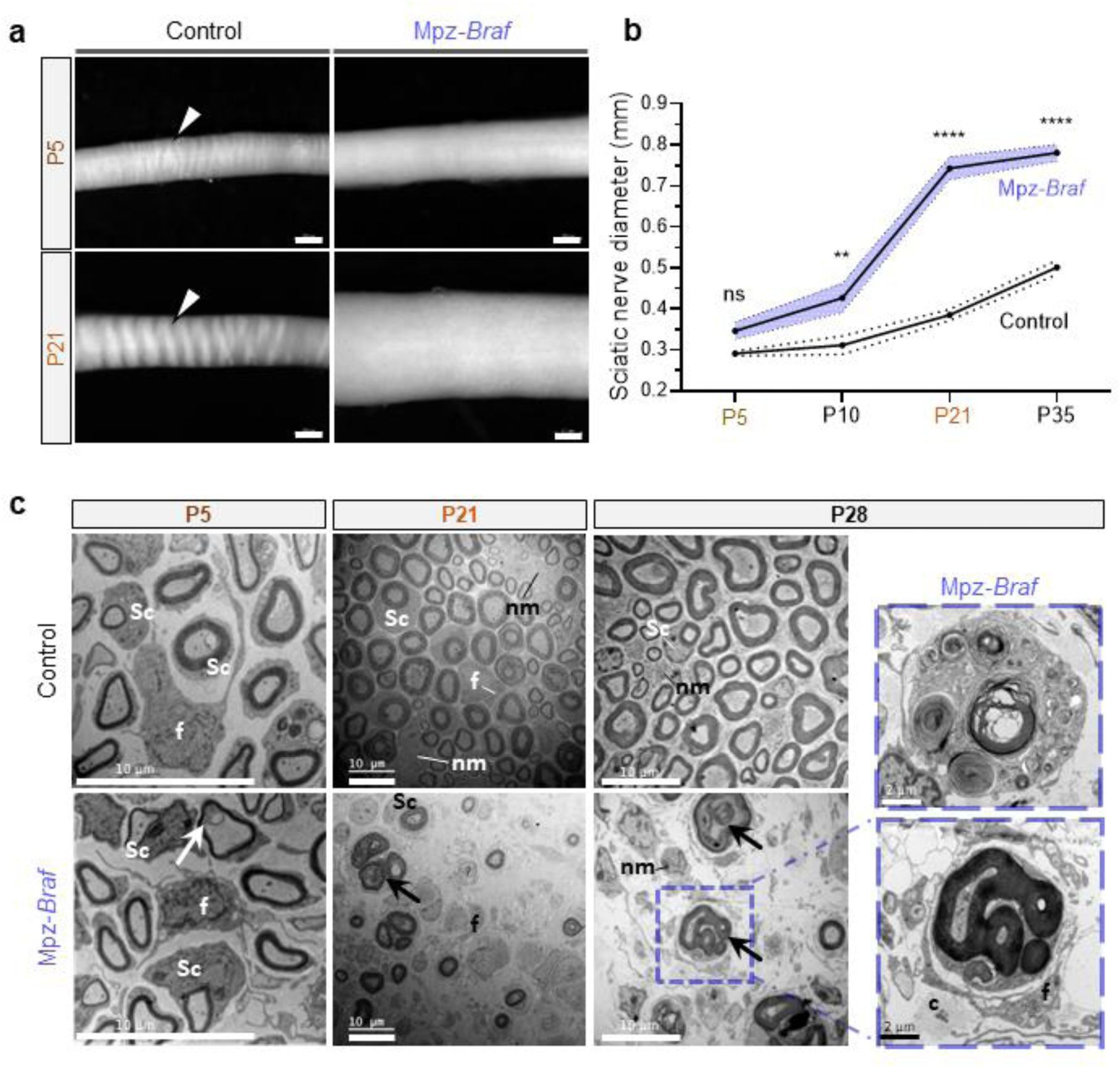
Myelination initiation is not impaired in *Braf*-mutant sciatic nerves, while maintenance is compromised. **a)** Macroscopic examination of control and mutant sciatic nerve at P5 and P21. Arrows indicate Fontana bands, present only in control nerves. **b)** Sciatic nerve diameter measurements at P5, P10, P21, and P35 in both Mpz-*Braf* and control mice (n = 11 *vs.* 6 at P5, n = 6 *vs.* 4 at P10, n = 15 *vs.* 10 at P21, and n = 5 *vs.* 5 at P35; in control *vs.* mutant nerves, respectively). Two-way ANOVA followed by Šidák’s post-hoc test; ns, not significant; ** *p* < 0.01; **** *p* < 0.0001. **c)** Representative transmission electron micrographs of sciatic nerve cross-sections at P5 and P21 from control and mutant mice. Bars = 10 µm except for magnified axons at P28, 2 µm. Arrows, myelin aberrations; c, extracellular collagen; f, fibroblast; nm, non-myelinating Schwann cell; sc, myelinating Schwann cell;.

To assess myelination dynamics and ultrastructure, we analyzed mutant sciatic nerves cross-sections using transmission electron microscopy (EM) at P5, P21 and P28. Over time, mutant nerves exhibited degenerating axons, extracellular collagen, fibroblasts, macrophages and abundant SC nuclei. By P21, myelin aberrations, including tomaculae, infoldings, onion bulbs degenerated axons, and focal hypermyelination of small-caliber axons, were frequent (**Fig 2c**; **Supplemental Figure S2a-d**). These anomalies were interspersed with abundant extracellular collagen fibrils and tentacular fibroblasts (**Fig 2c**). By P42, axon-associated macrophages containing myelin debris were also evident (**Supplemental Figure S2e-f**).

At P5, no significant differences were observed in average axonal density (52 ± 2 axons/mm^2^ in Mpz-*Braf vs.* 48 ± 3 axons/mm^2^ in controls) or diameter (1.447 ± 0.026 µm *vs.* 1.476 ± 0.072 µm; **Fig 3a**; **Supplemental Figure S3a-b**). Likewise, immune infiltration was absent, and myelin thickness as quantified by the *g*-ratio, inner/outer axonal myelin daimeter) ^40^, remained unchanged (0.732 ± 0.019 in mutants *vs.* 0.708 ± 0.028 in controls; **Fig 3b** and **Supplemental Figure S3c-d**). Radial sorting and myelination initiated similarly in mutants and controls, with only occasional myelin aberrations at this stage (**Fig 2c**).

**Figure 3.**
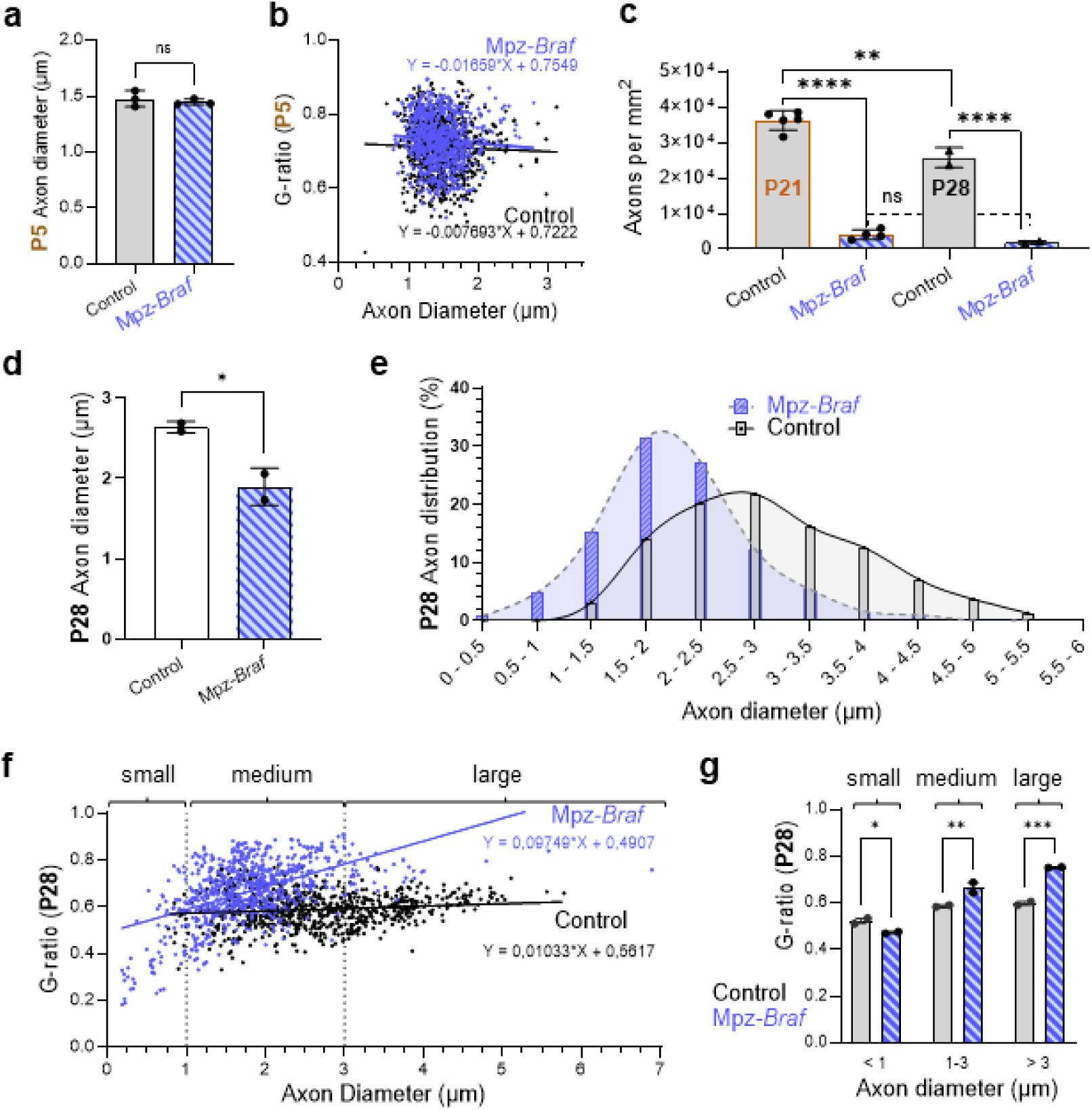
Braf^V600E^ in Schwann cells triggers demyelination and degeneration of large-caliber axons. **a)** Mean axon diameter at P5 (n = 3 per genotype, not significantly [ns] different). **b**) MyelTracer-obtained *g*-ratios as a function of axon diameter at P5, to assess myelin thickness in control (n = 803) *vs.* mutant (n=811) myelinated fibers (n=3 per genotype). The slopes of a simple linear regression do not vary significantly across P5 genotypes. **c)** Axon density decreases physiologically between P21 and P28 in control sciatic nerves but is significantly reduced to 7-11% of normal levels in mutant nerves at these stages. **d**) Mean axon diameter per genotype at P28 (n = 2 per genotype) shows that smaller fibers remain on average. **d)** Binning analysis of *g*-ratios as a function of axon diameter at P28. A total of n = 792 *vs.* 746 myelinated axons were analyzed for control *vs.* mutant nerves, respectively, from 2 individuals of each group (correlation significance indicated by asterisks). **f)** *g*-ratios of axonal diameters at P28, with a significantly different slope (p<0.0001) after simple linear regression, where mutant *g*-ratios increase most with size. **g**) When binned by diameter ranges (small 0-1 µm, medium 1-3 µm, and large >3 µm), there is modestly significant hypermyelination in small axons but more significant hypomyelination at diameters over 1 µm. Statistical significance was assessed with two-way ANOVA followed by Šidák’s post-hoc test; ns, not significant; * *p* < 0.05; ** *p* < 0.01; *** *p* < 0.001; **** *p* < 0.0001.

By P21, axon density had declined dramatically in Mpz-*Braf* mutants (4,060 ± 1,378 axons/mm^2^ *vs.* 36,172 ± 2,726 in controls; **Fig 3c**). At P28, only 7.2% of myelinated axons remained in mutant sciatic nerves (1,841 ± 465 axons/mm^2^ *vs.* 25,703 ± 2,812 axons/mm^2^ in controls; **Fig 3c**). Surviving mutant axons were significantly smaller (1.89 ± 0.23 µm *vs*. 2.64 ± 0.07 µm in controls; **Fig 3d-e**). To determine which axon calibers were affected, we analyzed 792 mutant and 746 myelinated axons at P28 (**Fig 3f**). Large-diameter axons (>3 µm) were disproportionately lost post-weaning (8.7 ± 6.7% in Mpz-*Braf vs*. 33.3 ± 5.6% in controls; **Fig 3e-f; Supplemental Figure S3e**).

Residual axons exhibited 12% higher *g*-ratios, indicating thinner myelin sheaths (*g*-ratios: 0.66 ± 0.02 in mutants vs. 0.59 ± 0.00 in controls; **Fig 3f; Supplemental Figure S3f**). When binned by caliber into small (<1 µm), medium (1-3 µm) and large (3 µm) diameters, medium- and large-caliber axons showed significantly thinner myelin in mutants (*g*-ratios: 0.67 ± 0.03 *vs.* 0.59 ± 0.01 for medium; 0.75 ± 0.00 *vs.* 0.60 ± 0.01 for large; **Fig 3g**). In controls, myelin thickness increases with axon size to plateau at axon diameters ≥4 µm ^30^. Conversely, small-caliber axons exhibited a modest (10.6%) but significant increase in myelination (*g*-ratios: 0.47 ± 0.01 in mutants *vs.* 0.52 ± 0.02 in controls; **Fig 2c**; **Supplemental Figure S2d**). Fast-conduction motoneuron axons ^41^ were thus most severely affected in mutants, while small axons were often hypermyelinated.

### Myelin component mislocalization and SC progenitor expansion

To extend our structural findings, we examined myelinated axon components by teasing fixed fibers from P5 and P21 sciatic nerves. Immunofluorescence for contactin-associated protein 1 (Caspr/paranodin) revealed that, unlike controls (**Fig 4a**), mutant axons displayed irregularly spaced, diffuse or absent myelin internodes (**Fig 4b, d-g**). At P5, longitudinal views already showed early myelin breakdown, with F-actin labeling highlighting beaded myelin ovoids along some mutant fibers (**Fig 4b-d**). By P21 (**Fig 4e**), Mpz-*Braf* mice alone displayed demyelinated segments lacking myelin basic protein (Mbp) (**Fig 4f**). The presence of multiple adherent nuclei between residual Caspr+ nodes (**Fig 4f**), rather than the single SC expected after normal radial sorting ^42^, suggests persistent immature SCs in mutants.

**Figure 4.**
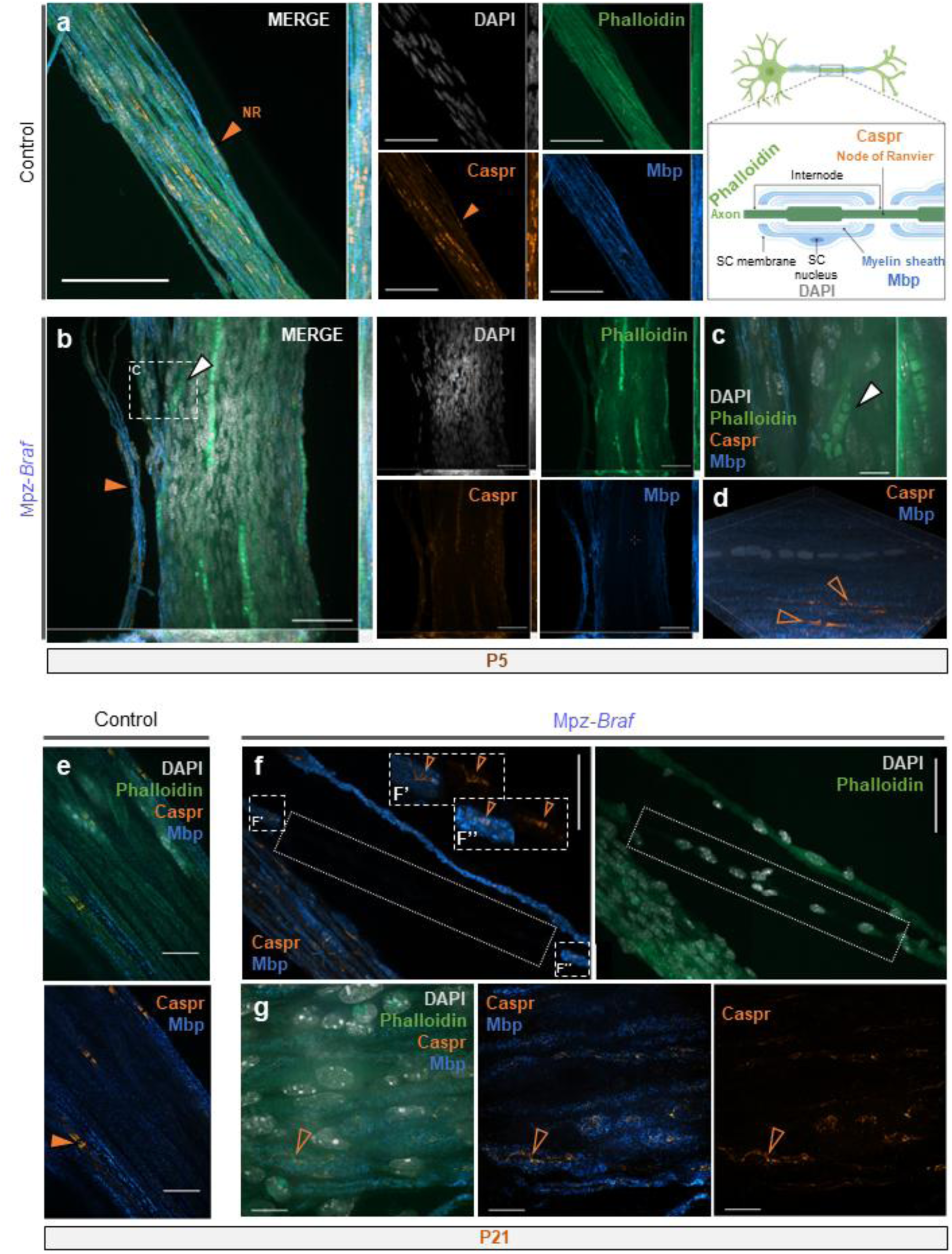
Segmental demyelination is preceded by myelin disassembly in Mpz-*Braf* mice. Teased fibers from sciatic nerves of control (**a, e**) and Mpz-*Braf* mice (**b-d**, **f-g**) at P5 (top panels) and P21 (bottom panels) were immunostained with antibodies against Mbp and Caspr and Alexa Fluor-488-conjugated phalloidin. Mpz-*Braf* fibers had degenerating myelin ovoids (white arrowheads), fewer apparent nodes of Ranvier with diffusing paranodal Caspr (orange arrowheads), and segmental demyelination, not observed in controls. Nuclear DNA was stained with DAPI (grey). Maximum intensity z-projection images were generated in **b, e** and **f** with Nikon NIS-Elements Viewer software. Scale bars: 50 µm (**a, b, f**) or 10 µm (**c, e, g**).

To assess proliferating SCs, we examined the co-expression of Ki67, a cell cycle progression marker ^43,44^, and Sox10 (SRY-box transcription factor 10), present throughout the SC lifespan ^45^, by immunostaining sciatic nerve sagittal sections at P42. While control nerves showed sparse Sox10^+^/Ki67^+^ nuclei (**Supplemental Figure S4a**), mutant hyperplastic nerves exhibited densely, irregularly arranged double-positive nuclei (**Fig. S4b**). Notably, all mutant Sox10^+^ nuclei were strongly Ki67^+^ in mutant nerves, interspersed with additional Ki67^+^ Sox10^-^ cells. These findings indicate nerve hyperplasia rather than hypertrophy in mutants, as sustained SC proliferation is concomitant with inflammation and fibrosis.

### *Braf*^V600E/+^ glia *in vivo* downregulate myelin sheath gene transcription

To identify global transcriptional changes associated with myelin loss after MAPK pathway activation, we performed bulk RNAseq on Mpz-*Braf* and control sciatic nerves dissected from four mice each at P5 and P21 (**Fig 5** and **Supplemental Table S2**). Unlike P5 pituitaries, 2,540 differentially expressed genes (DEGs; padj< 0.05) were identified at P5 between mutant and control nerves. By P21, this rose to 10,409 DEGs (**Fig 5a**), indicating early, large-scale transcriptional changes in sciatic nerve cells following constitutive MAPK signaling.

**Figure 5.**
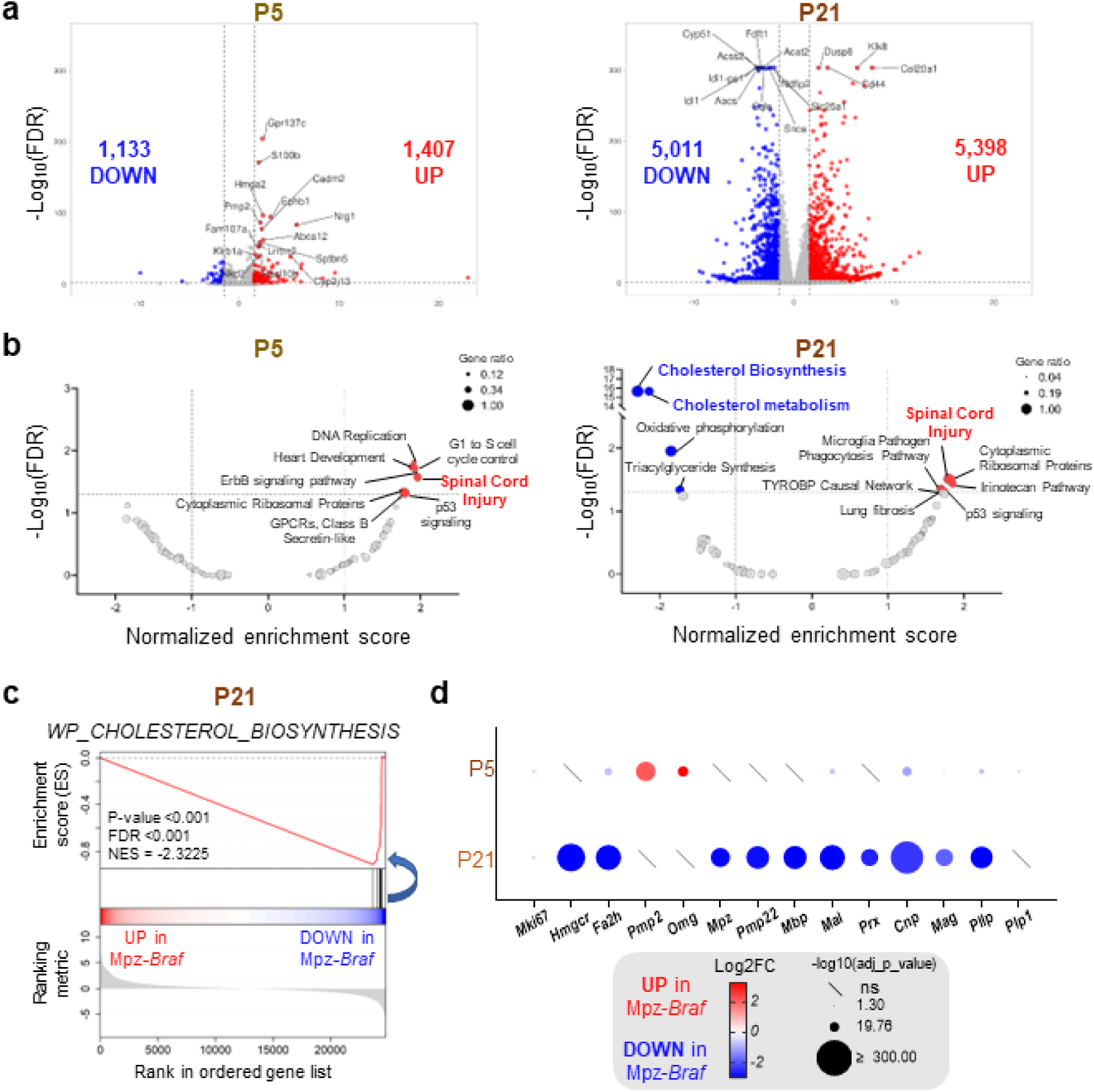
Comparative transcriptomics show downregulation of myelin sheath component transcripts in *Braf*-mutant sciatic nerves. **a)** RNA-seq analysis was performed in control and Mpz-*Braf* mutant sciatic nerves (n = 4 per genotype). Volcano plots depict the DEGs in *Braf*-mutant sciatic nerves relative to controls at P5 (left panel) and P21 (right panel). An adjusted *p*-value < 0.5 and Log2FC <-1 (blue) or >1 (red) were used as thresholds to define DEGs. The top 15 DEGs are labeled on each graph according to Manhattan distance. **b)** Volcano plots illustrate the differentially enriched pathways provided by GSEA through WebGestalt https://www.webgestalt.org/ ^117^ using the mouse Wikipathway database. Pathways are significantly enriched when false discovery rate (FDR) < 0.05 and normalized enrichment score (NES) < -1 (blue) or >1 (red). Circle diameter is proportional to gene ratio (count of core enrichment genes / count of pathway genes). **c)** Enrichment plot for “cholesterol biosynthesis”, the most significant gene set downregulated in mutant sciatic nerves (p-value and FDR < 0.001, NES = -2.3). **d)** Dot plot displaying log2 fold-change (Log2FC) of myelin-associated genes (*Mpz*, *Pmp22*, and *Mbp* in bold, representing transcripts of the primary myelin sheath proteins).

The myelin sheath comprises 70% to 85% lipids and 15 to 30% proteins ^46^. Notably, the term “spinal cord injury” was enriched in DEGs at both P5 and P21 (**Fig 5b**). By P21, cholesterol biosynthesis and metabolism pathways were significantly downregulated in mutant nerves (**Fig 5b, c**). Strongly downregulated DEGs at P21 included *Hmgcr* (11.5-fold reduction), encoding the rate-limiting enzyme in cholesterol biosynthesis ^47^, and *Fa2h* (7.9-fold reduction), encoding fatty acid 2-hydroxylase, a key enzyme in myelin glycosphingolipid synthesis ^48^ (**Supplemental Table S2**). While *Pmp2* (Fabp8, a fatty acid-binding protein) and *Omg* (oligodendrocyte myelin glycoprotein) were significantly upregulated at P5 in mutants, their expression normalized to control levels by P21 (**Fig 5d**). Transcripts for other myelin proteins (e.g., *Mpz*, *Pmp22* [peripheral myelin protein 22], *Mbp* [myelin basic protein]) showed no significant changes at P5, but other myelin-associated transcripts, including myelin and lymphocyte protein (*Mal*), periaxin (*Prx*), the myelin-associated 2’,3’-cyclic nucleotide 3’ phosphodiesterase (*Cnp*), myelin-associated glycoprotein (*Mag*), and plasmolipin (*Pllp*) were already modestly but significantly reduced **(Fig 5d**; **Supplemental Table S1**). By P21, when control sciatic myelination is nearly complete (**Fig 2**), *Mpz*, *Mbp* and *Pmp22* transcripts were also significantly downregulated in mutants (**Fig 5d**). Thus, sustained MAPK signaling drives transcriptional downregulation of myelin components, contributing to the observed demyelination and axonal degeneration phenotype.

### Constitutive MAPK signaling disrupts transcriptional control of myelination

To investigate how constitutively active MAPK signaling modulates myelination-associated transcription factors (TFs), we analyzed co-variance patterns in Mpz-*Braf* and control sciatic nerves. At P5, key myelination activators *Egr2* (Krox20; early growth factor response 2 ^49^), and *Sox10* ^50^, remained unchanged. However, by P21, mutant SCs exhibited a significant decrease in Egr2 and the melanocyte determinant *Mitf*, with concomitant ectopic increases in neural crest or multipotency maintenance TFs *Sox2, Foxd3*, *Tfap2a* and *Tfap2b,* with no change in *Sox10* (**Fig 6a**; **Supplemental Table S2**). Concurrently, myelination transcriptional repressors ^51^ were significantly upregulated in mutant nerves at P21, including *Jun*, *Egr1*, *Egr3,* and *Zeb2* (**Fig 6a**). Increased *Jun* translated to higher Jun protein levels in mutants beyond P21, whereas Egr2 protein loss became detectable at P42 (**Fig 6b**).

**Figure 6.**
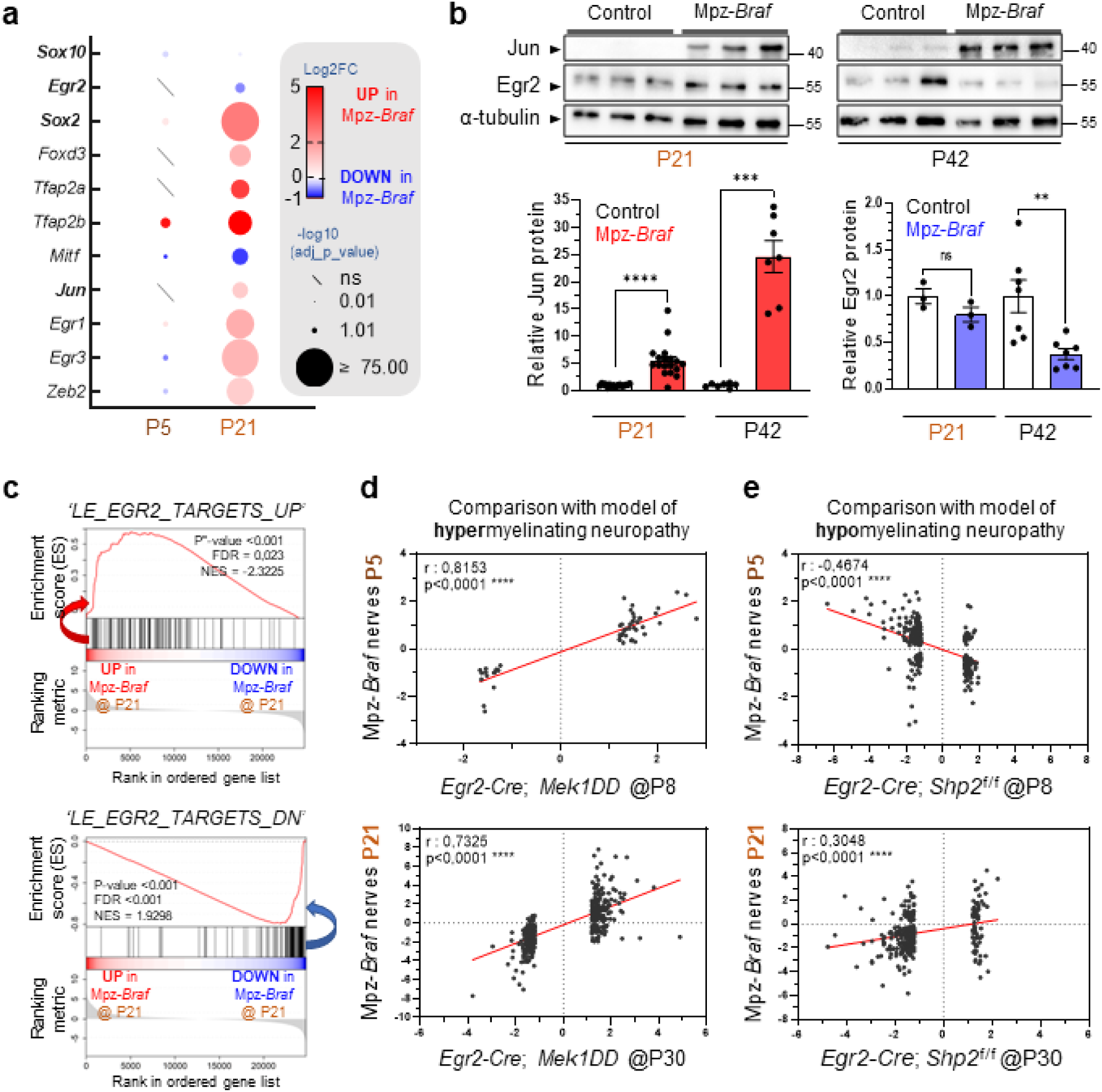
*Egr2* and its target genes decrease in Mpz-*Braf* nerves with concomitant increase in expression of negative regulators of myelination. **a)** Dot plot displaying log2 fold-change (Log2FC) of known positive (*Sox10*, *Egr2*) or negative (*Jun*, *Sox2*) transcriptional regulators of myelination in RNAseq of mutant *vs.* control sciatic nerves at P5 and P21. **b)** Western blot of Jun and Egr2 in sciatic nerve lysates from P21 and P42 Mpz-*Braf* and control mice. Gapdh was detected as a loading control. N=13 individuals of each genotype. Statistical significance was assessed by an unpaired Mann-Whitney test; ns, not significant; ** *p* < 0.01; *** *p* < 0.001; **** *p* < 0.0001. **c)** Enrichment plot for up- or down-regulated genes (top and bottom panels, respectively) showing that most genes are commonly deregulated in sciatic nerves from either the *Mpz-Braf* or homozygous *Egr2* hypomorphic allele mice ^132^. **d-e)** Spearman’s *rho* to assess significance of correlation was calculated from bulk RNAseq of Mpz-*Braf* sciatic nerves at P5 (top panels) or P21 (bottom panels), with analogous datasets at P8 and P30 from Egr2-*Cre*-driven (**d**) overexpression of constitutively active Mek1 (Egr2*-Cre*; *Mek1*DD) or (**e**) knockout of *Shp2* (Egr2*-Cre*; *Shp2*^f/f^).

To corroborate these findings, we compared our model to a hypomyelinating neuropathy model with germline expression of two hypomorphic *Egr2* alleles (Le et al., 2005). Gene Set Enrichment Analysis (GSEA) analysis revealed significant co-variance of 90% (99/110) upregulated and 91.5% (98/107) downregulated DEGs (**Fig 6c**; *p* < 0.001). Sciatic nerve hypomyelination is therefore linked to increased Jun expression, followed by reduced *Egr2*-driven transcription.

During remyelination after sciatic nerve injury, axonal Nrg1 type III stimulates SC *Pmp2* expression, facilitating fatty acid uptake and mitochondrial ATP production ^52^. In Mpz-*Braf* mice, RNAseq revealed significant *Pmp2* and *Nrg1* overexpression (**Supplemental Table S2**). Notably, the major source of Nrg1 after P5 is the SC rather than the axon itself ^53^. This reactive overexpression to induced injury or potentially physiological wear, associated with “onion bulb” formation around neuropathic axons, contributes to an ultimately detrimental chronic repair response ^54^.

Nrg1-III normally binds SC-specific ErbB3 to activate MAPK signaling. We hypothesized that Mpz-*Braf* sciatic nerves (P5/P21) should mirror the transcriptional response of an *Egr2*-driven gain-of-function *Mek1*DD allele (P8/P30), which bypasses the natural Erbb3 decline in maturing SC and hypermyelinates ^30^. Since *Mek1* functions downstream of *Braf*, we observed robust positive correlation (*p* < 0.0001) at both myelination stages as well as significant *Erbb3* upregulation between P5 and P21 **(Fig 6d**; **Supplemental Table S2)**.

Conversely, *Egr2*-Cre-driven Shp2 depletion (encoded by *Ptpn11*, a Noonan syndrome RASopathy gene ^55^) causes hypomyelinating neuropathy ^30^. Comparing Mpz-*Braf* and *Egr2^Cre^;Shp2^f/f^*sciatic transcriptomes revealed significant anti-correlation of DEG at initial myelinization (P5 vs. P8) but positive correlation post-weaning (P21 vs. P30; **Fig 6e**).

Finally, to test whether oncogenic Braf effects could be recapitulated in *Egr2*-expressing (rather than *Mpz*-expressing) SCs *in vivo*, we crossed heterozygous LSL-*Braf*^V600E/+^ mice with the same *Egr2*-Cre drivers. The *Egr2*-Cre allele is expressed by SCs from mid-gestation on (Topilko et al., 1994; **Supplemental Figure S5A**). However, no mutant neonates survived, deviating significantly (*p* < 10^-6^) from expected Mendelian ratios. This fully penetrant embryonic lethality, likely due to broader expression of *Egr2* beyond SCs ^56,57^, prevented further postnatal myelination comparisons (**Supplemental Figure S5B**).

### Uninjured sciatic nerves initiate and sustain reprogramming to an early repair program

Transcripts significantly overexpressed at P5 and sustained through P21 may reflect direct rather than indirect effects of MAPK signaling. We therefore examined 175 genes that were significantly upregulated by ≥2-fold in both P5 and P21 mutant sciatic nerves. Pathway enrichment analysis (Wikipathways, KEGG, REACTOME (**Supplemental Figure S6a**) and GSEA (**Supplemental Figure S6b**; **Fig. 7a**) revealed that “spinal cord injury” (WP2431) remained one of the most significantly enriched terms (cf. **Fig 5b**). Notably, cell adhesion molecule L1 (*Chl1*), a highly specific repair-type SC marker identified in single-cell RNAseq ^23^, was significantly enriched. Comparing the Mpz-*Braf* sciatic nerve transcriptome to a 6- to 8-week mouse axotomy model at seven days post-injury ^58^ showed significant positive correlation at P5 (**Supplemental Figure S6c**), which strengthened at P21 (**Fig 7b**). Following physical injury (e.g., crush or axotomy), SCs re-express immaturity genes and transcribe novel repair-associated genes. Jun, a repair SC-specific TF that represses myelination by opposing Egr2 ^59^, was significantly more transcribed and translated at P21 in the Mpz-*Braf* sciatic nerve than controls (**Fig 6a-b**). Thus, juvenile Mpz-*Braf* mice misactivate a molecular repair response during physiological myelination.

**Figure 7.**
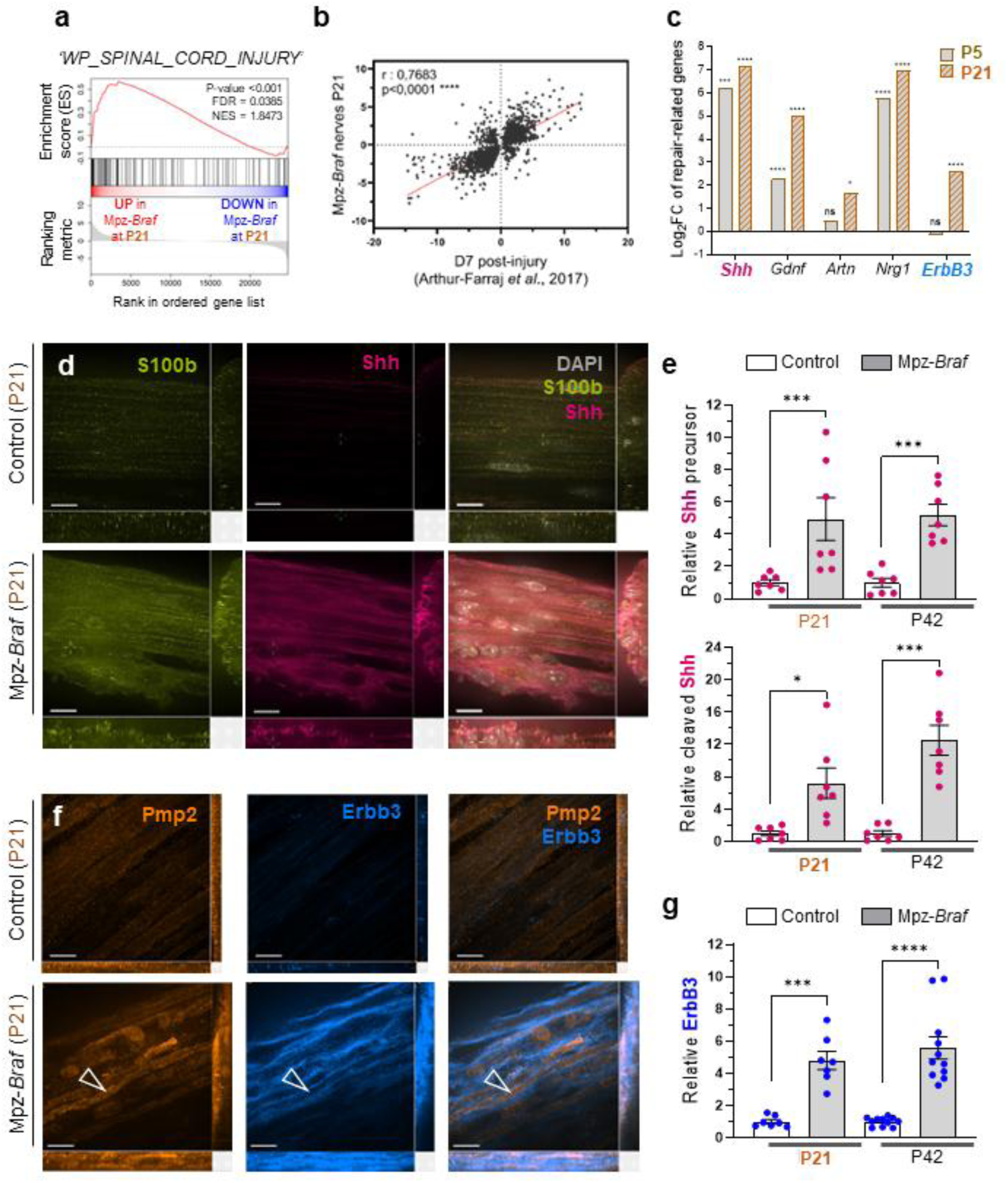
Activation of injury-related genes in Mpz-*Braf* sciatic nerves. **a**) GSEA analysis of upregulated genes at P21 shows highly significant correlation with the Wikipathway annotation "Spinal cord injury" (WP2431). **b**) These are also significantly and positively correlated with genes upregulated in regenerating nerves of young adult mice 7 days after axotomy. **c**) Transcription of *Shh*, *Gdnf* and *Nrg1* are greatly increased at both P5 and P21, while Gdnf-family member *Artn* and *Nrg1* receptor ErbB3 are only significantly upregulated at P21. Teased fibers from sciatic nerves of P21 control and Mpz-*Braf* mice immunostained against (**d**) S100b (yellow) and Shh (magenta), with DAPI nuclear counterstain (gray), or (**f**) Pmp2 (orange) and Erbb3 (blue). Scale bars: 50 µm (**d**) or 10 µm **(f**).

To determine whether Jun-dependent repair ligand effectors were also upregulated, we examined *Shh* (Sonic hedgehog) and the neurotrophic factor *Gdnf* (glial cell line-derived neurotrophic factor) ^60^. Both were significantly upregulated as early as P5 (**Fig 7c**). Cleaved Shh, critical for multipotent NC maintenance ^61^ and reactivated in repair SCs ^62^ was confirmed by immunolabeling of fiber bundles (**Fig 7d**) and Western blot **(Fig. 7e**; **Supplemental Figure S7e**). S100b, a calcium-binding intermediate filament in SCs and their multipotent progenitors ^63^, was also co-localized and significantly overexpressed by P21 in mutants.

Additionally, Nrg1-III and the Gdnf family member Artemin (Artn), secreted by Jun-positive repair SCs to promote axon outgrowth ^53,64^, were upregulated at P5 (*Nrg1*) and P21 (*Nrg1*, *Artn*) (**Fig. 7c**). The Nrg1 receptor *ErbB3*, a nerve damage hallmark ^65^, was transcriptionally and translationally upregulated over time (**Fig 7c, f-g**; **Supplemental Figure S7b**), with increased Erbb3 immunofluorescence in mutants. This distribution was complementary to Pmp2-positive myelin in individual cells (**Fig. 7f**; **Supplemental Figure S7d**). Vimentin (*Vim*), an intermediate filament upregulated in injured axons ^66^, also increased over time (**Fig. S7c; Supplemental Table S2**). Despite repair program activation, progressive axonal degeneration in mutants implies that sustained MAPK signaling is counterproductive for SC-mediated trophic activity ^67^ *in vivo*.

### Schwann cell differentiation of CFC patient-derived stem cells is impaired

While the Mpz-*Braf* model demonstrated phenotypic consequences of MAPK overactivation, germline *BRAF*^V600E^ is embryonic lethal. To study a RASopathy-associated variant ^68^, we differentiated human induced pluripotent stem cells (hiPSCs) from two unrelated CFC patients with heterozygous *BRAF*^Q257R^ variants ^68,69^ into SCs *in vitro* ^70^ and compared to two unrelated control lines. Control hiPSCs followed expected differentiation ^70^, but one mutant line (RMK0138C) required more frequent passaging to maintain similar density and failed to develop Schwann cell progenitor (SCP)-like morphology (elongated, refringent soma and long projections) even after 90 days (**Fig 8a**).

**Figure 8.**
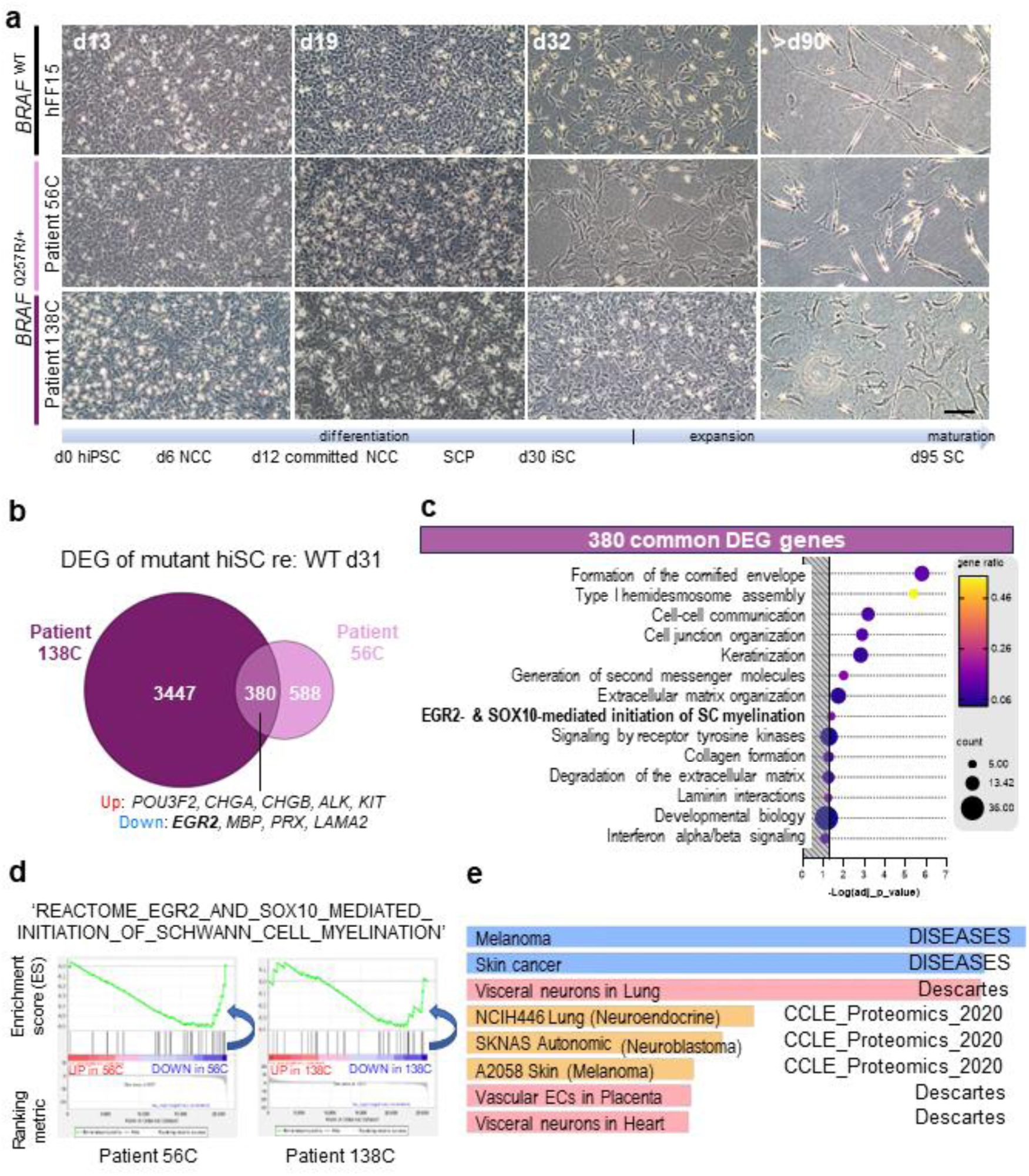
Patient-derived immature Schwann cells express inappropriate gene signatures *in vitro*. **a)** Summary of hiPSC-directed differentiation into SC, derived from two unrelated WT (HFF15 shown here and AG08H) and from unrelated adolescent girls, both carrying the *BRAF* p.Q257R variant (CFC patients RMK0056C, RMK0138C) over time. Representative bright-field images shown at D13, D19, a day after passage on D32, and at D95-100. Scale bar = 100 µm for all frames. **b**) Euler diagram showing the number of common/specific dysregulated genes (*p*-value < 0.05, Log2FC > 1 or < -1) in RMK0056C and RMK0138C mutant cells compared to control cells (HFF15) at D21 of differentiation. (C) Bubble plot of enrichment analysis of the common 380 dysregulated genes in RMK0056C and RMK0138C. Gene set names from the REACTOME database are indicated on the y-axis. Gene sets are ranked according to significance level (-log adjusted p-value). The size of the dots represents the number of queried genes present in the corresponding pathway, and the color corresponds to the gene ratio (number of queried genes present in the gene set / gene set size). (D) Enrichment plot for Egr2- and Sox10-mediated initiation of SC myelination using Reactome database, corresponding to the significant gene set downregulated in RMK0056C and RMK0138C cell lines after 30 days of differentiation. (E) Enrichr-KG analysis of the most significantly enriched terms across three additional databases (Descartes Cell Types and Tissue 2021, Jensen DISEASES and CCLE Proteomics 2020) highlight the close lineage relationship between Schwann cell progenitors and either melanocytes (“melanoma” and “skin cancer”, the A2058 cell line) or neuroendocrine cells (“visceral neurons in lung/heart”, the SKNAS and NCIH446 cell lines).

RNAseq at (d)18, a committed SCP-like stage, and d31, analogous to immature SC (**Fig 8a**) reveealed 380 DEGs in both mutant lines (**Supplemental Table S3**). Reactome ^71^ pathway analysis highlighted their unexpected downregulation of “epidermal cornification” (REAC:R-HSA-6809371), including eight keratin genes (**Fig 8c**; **Supplemental Table S3** and suppression of “EGR2 and SOX10-mediated initiation of SC myelination” (REAC:R-HSA-9619665). Downregulated transcripts included *EGR2* itself*, MBP*, *PRX* and *LAMA2*, encoding the alpha-2 chain of laminin-211, a component of SC basal lamina necessary to restrain small-caliber fiber hypermyelination ^72^ (**Fig 8c-d)**.

In contrast, *POU3F2*, a TF transiently required in neonatal murine SC ^73^, was significantly upregulated in both CFC patient-derived cell lines (**Supplemental Table S3**). DEGs co-expressed with *POU3F2* evoked multipotency of neural crest-derived SCPs. Upregulated annotations included “melanoma” and the proteome of amelanotic, *BRAF*^V600E^-mutated melanoma cells (A2058); proteomes of *P53*-mutated lung neuroendocrine carcinoma (NCI-H446) and *NRAS*^Q61L^-mutated neuroblastoma (SK-N-AS) cells, and transcripts common to “visceral” (parasympathetic) autonomic neurons **(Fig 8e**). These findings suggest that prolonged postnatal MAPK signaling in immature SC, both in mice and humans, sustains transcriptional programs typical of other neural crest lineages ^28,74^. Thus, persistent MAPK pathway activation in cycling SC prevents their terminal differentiation *in vitro* and *in vivo*.

## Discussion

### Hyperplastic neuropathy as a shared RASopathy feature

Our findings demonstrate that constitutive activation of BRAF-mediated signaling in a subpopulation of NC-derived cells recapitulates systemic effects found across multiple germline RASopathy syndromes. A key phenotype was rapid onset of hyperplastic neuropathy, reminiscent of “hypertrophic neuropathy” described in case reports ^18,20,75–78^. This phenotype had not been reported in prior systemic RASopathy animal models ^79^, suggesting that neuropathy should be systematically assessed in RASopathy patients, given the mechanistic convergence on sustained ERK1/2 activation in Schwann cells.

### Growth retardation and endocrine dysregulation

Like other RASopathy models (e.g. germline mutations in *Kras* ^80^, *Braf* ^81–84^, *Raf1* ^85^, *Ptpn11* ^86^ or *Sos1* ^87^), the Mpz-*Braf* mouse presents early postnatal failure to thrive. Severe feeding difficulties are common in *BRAF*-associated germline RASopathies ^18,88–90^. However, Mpz-*Braf* mice exhibited normal feeding behavior despite motility issues, while growth delay due to Igf1 deficiency preceded weaning. No megacolon was observed at any age as a sign of gut dysmotility, but transcripts co-expressed specifically in adult enteric glia, such as *Plp1* and *Gfap* ^91^, were upregulated over time in Mpz-*Braf* SC (**Supplemental Table S3**). This suggests the repair phenotype reverts SCPs to a commitment state preceding glial specialization.

By restricting strong MAPK activation to *Mpz*-expressing cells, including SCs and SCPs ^92^, we unintentionally also targeted a subset of Gh-producing pituitary cells (somatotrophs) that can express *Mpz* under conditions of systemic inflammation (https://cells.ucsc.edu/?ds=mouse-pituitary-inflam&gene=Mpz) ^93^. Postnatal *Igf1*-null mice, which survive at low rates, exhibit severe growth retardation ^94,95^, while liver-specific Igf1 rescue restores both body size and fertility ^96^. The fully penetrant cryptorchidism in male Mpz-*Braf* but not *Igf1*-null mice, despite no *Mpz* expression in germ cells, suggests a neurogenic mechanism. Cryptorchidism is common in RASopathies ^4,97^, and genitofemoral nerve damage in rats causes testicular ascent^98^. This supports a role for peripheral nerve myelination in maintaining testicular descent and fertility ^99,100^. Uncoupling of Gh-Igf1 by sustained MAPK signaling mirrors the effects of blocking differentiation of both SC (*via* Nrg1-ErbB3) and pituitary progenitors (eg. *Prop1*-driven *Braf* models) ^2^, implicating constitutive MAPK activation in additional human disorders.

### Future directions in congenital hypomyelinating neuropathy and lineage plasticity

Severe, early-onset CMTs, caused by mutations in *MPZ*, *EGR2*, *PMP22* or *PRX* ^22^, share features with our model: demyelination, profuse SC proliferation, distal sensory loss, and the enlarged, “gelatinous” peripheral nerves first reported in 1893 ^101^, which principally affect the legs. Mpz-*Braf* mice exhibited hindlimb tremor and locomotor deficits, extending to other body areas by P42 but with generalized nerve enlargement by P21. Our findings confirm that SCs initiate these neuromuscular outcomes. The absence of apparent suffering (e.g., aversion, unkempt fur, flattened ears), makes the weanling model tractable for behavioral or electrophysiological studies of RASopathy-associated neuropathies ^102^.

Furthermore, patient-derived, *BRAF*^Q257R^-expressing NC cells are stochastically disadvantaged in adopting a differentiated SC phenotype *in vitro*, as compared to *BRAF*-wildtype cells. Prospective nerve biopsies in unresolved cases of either RASopathies or early-onset CMT could confirm these findings, linking transcriptional dysregulation to clinical phenotypes and improving molecular diagnoses while opening new interventional avenues.

## Methods

### Ethical statement

This study complies with all relevant ethical regulations. Animal care and experiments were in accordance with French and European legislation and approved by the French ACUC (C2EA-14; project numbers 37-08102012 and 9522-2017040517496865v5). Human iPSC studies adhered to the Declaration of Helsinki ^103^ and French bioethics law (2021-1017)^104^ consistent with our institutional review board protocols.

### Mouse breeding and genotyping

Transgenic *P0-Cre^+/°^*; *Braf*^V600E/+^ and *Krox20-Cre^+/°^*; *Braf*^V600E/+^ mice were generated by crossing *LSL-Braf*^V600E/+^ mice ^11^ (RRID:IMSR_JAX:017928) with P0(Mpz)-*Cre* (RRID:IMSR_JAX:017927) ^36,105^ or Krox20-*Cre* (RRID:IMSR_JAX:025744) ^106^ drivers. Genotyping was performed by PCR (see **Supplemental Methods**). Mice were housed under standard conditions (12-hour light/dark cycles, *ad libitum* access to food/water).

### Western Blots

Sciatic nerve lysates were prepared in RIPA buffer with protease inhibitors, separated by 12% SDS-PAGE, and probed with antibodies against phospho-Erk1/2 (RRID:AB_331646), Erk1/2 (RRID:AB_330744), c-Jun (RRID:AB_2130165) and Gapdh (RRID:AB_2107445). Alternately, primary antibodies to Shh (RRID:AB_528466)^107^, Erbb3 (RRID:AB_358277), Egr2 (RRID:AB_10862073), alpha-tubulin (RRID:AB_2241126), vimentin (RRID:AB_2881439) or vinculin (RRID:AB_10603627) were used. Signals were detected using relevant secondary antibodies (RRID:AB_330924 and RRID:AB_2099233) followed by chemiluminescence detection and quantified with ImageJ v.1.54f.

### Culture and differentiation of human pluripotent stem cells (hiPSC)

Two female patients with cardio-facio-cutaneous (CFC) syndrome bearing a heterozygous germline *BRAF* Q257R variant, or their legal guardians, provided written consent as approved by the UCSF Human Research Protection Program (CHR #10-02794), to produce the human induced pluripotent stem cell (hiPSC) lines, RMK0056C and RMK0138C, as described ^69^. The AG08H and HFF15 clones of healthy donor male hiPSC were derived from commercial cell lines consented for research, AG08498 (RRID:CVCL_1Y51) and HFF-1 SCRC-1041 (RRID:CVCL_3285) as described ^108^. Clones were initially grown in mTeSR1 (Stemcell) on human ES cell-qualified Matrigel™ before cryopreservation. Thawed hiPSC were cultured on Synthemax II-SC (Corning) at 0.025 mg/mL and maintained with daily changes of StemMACS iPS-Brew XF (Miltenyi Biotec). At 60-80% confluence, they were passed with ReleSR (StemCell Technologies), with 10 μM Y-27632 for the first day. Differentiation of hiPSCs into SC ^70^ was initiated at 60% confluence and passed every 6 days until day 31. Thereafter, upon 90% confluence, they were split 1:2.

### RNA sequencing and analyses

Sciatic nerves and pituitary glands from WT or *Braf^V600E/+^*animals, and pellets of WT or *BRAF*^Q257R/+^ hiPSC-derived SCP and immature Schwann cells (iSC) were stored at -80°C. Tissues were lysed in the FastPrep-24 homogenizer (MP Biomedicals) and total RNA extracted with the RNeasy Lipid Tissue Mini (Qiagen) or, for cell pellets, NucleoSpin RNA Plus S (Macherey-Nagel) kits. RNA concentrations were measured on a NanoDrop TM 1000 Spectrophotometer (ThermoFisher). Libraries were prepared with the KAPA mRNA HyperPrep kit (Roche) and sequenced on a NovaSeq 6000 (Illumina). Quality was assessed using fastQC (v0.11.5). Raw sequencing reads were aligned to GRCm39/GRCh38 using STAR (v2.7.2b). Bam files were indexed and sorted with Samtools (v1.7). After mapping, reads per feature (Ensembl or GENCODE v34 annotations for mouse or human data, respectively) were determined using StringTie v2.1.6, and differential expression analyzed with a Wald test in DESeq2 (v1.34.0; FDR < 0.05). Data visualization made use of R packages Pheatmap and EnhancedVolcano ^109^. Expression-dependent enrichment analyses were conducted with Gene Set Enrichment Analysis (GSEA) v4.3.2 ^110^ using mouse Wikipathway and human Reactome databases, otherwise with g:Profiler ^111^ or Enrichr-KG ^112^, interrogating quantitative proteomics of the Cancer Cell Line Encyclopedia ^113^ KEGG, Reactome, Gene Ontology datasets.

These were significantly enriched by Enrichr-KG ^112^, using databases of annotations related to human disease ^114^, and a large single-cell atlas of developmental human gene expression ^115^.

### Nerve and fiber immunostaining

Sciatic nerves were fixed in 4% buffered paraformaldehyde and stained with primary antibodies to Mbp (RRID:AB_94975); Ki-67 (RRID:AB_10854564); Caspr (RRID:AB_869934); Sox10 (RRID:AB_2941085); Pmp2 (RRID:AB_2166978); Erbb3 (RRID:AB_358277); S100b (RRID:AB_1856538); or Shh (RRID:AB_528466)^107^, followed by secondary antibodies and DAPI counterstaining. Further details are in the **Supplemental Methods**. Image stacks of teased fibers were acquired with a spinning disk confocal microscope (Nikon SoRa) and NIS-Elements software and nerve images on a Zeiss LSM800 with Airyscan and Zen software v2.3.

### Transmission electron microscopy (TEM) and morphometry

Sciatic nerves were fixed in 2% PFA + 2.5% glutaraldehyde, post-fixed with 1% OsO_4_, contrasted in 1% uranyl acetate before Epon resin embedding. Semi-thin sections at 1 µm were counterstained with toluidine blue. Ultra-thin sections were imaged on an FEI Morgagni 120 kv microscope. Myelin morphological parameters were analyzed with MyelTracer ^40^. Axon density was assessed with ImageJ using the semi-thin sections.

### Statistics

Statistical analyses were performed with Prism v10.2.0 (GraphPad), using unpaired Mann-Whitney or two-way ANOVA non-parametric tests. For multiple comparisons, two-way ANOVA followed by Šidák’s post-hoc tests, or Student t-tests followed by Benjamini-Hochberg adjustment, were used. Mendelian segregation was assessed with a chi-square contingency test. Data are presented as mean ± SEM; significance: *\* p* < 0.05; ** *p* < 0.01; *** *p* < 0.001; **** *p* < 0.0001.

## Data availability

Raw and processed RNA-sequencing data from this study have been deposited to the Gene Expression Omnibus (GEO) and assigned the accession identifiers GSE262047 and GSE262048.

## Websites

"Catalogue Of Somatic Mutations In Cancer" (COSMIC), https://cancer.sanger.ac.uk/cosmic ^1^.

Rare disease ORPHA codes are maintained at https://www.orpha.net and available on https://www.orphadata.com/ ^116^.

fastQC (v0.11.5) is available at https://www.bioinformatics.babraham.ac.uk/projects/fastqc/ and EnhancedVolcano at https://bioconductor.org/packages/EnhancedVolcano.

NIH Fiji/ImageJ software (https://imagej.net/ij/) is available from https://imagej.net/ij/.

## Author contributions

E.M. wrote the first draft, performed and analyzed experiments; D.A. provided analyses, performed experiments and revised the article; P.Q., G.M. and M.M. performed experiments; N.B.M. and V.D. provided protocols and guidance; C.E.Y. and N.B. provided control hiPSC lines and guidance; L.A.W. provided patient hiPSC lines; A.B. provided constructive criticism and funding; H.C.E. supervised the experiments, provided funding, wrote and revised the article.

## Acknowledgements

We are grateful to C. Humbert, C. Castro, N. Trévisiol and C. Aubert of the MMG Genomics and Bioinformatics facility (GBiM) for assistance with RNAseq and analyses; Dr. F. Magdinier and O. Hadadeh of the Marseille Stem Cell (MaSC) core facility for control hiPSC cell lines and their quality control; Dr. T.-T. Mac, Adeline Querdray and Dr. T. Fauquier for their technical assistance and advice; the PiCSL-France BioImaging core facility, supported by ANR-10-INBS-04, and Dr. C. Leterrier and Dr. L. Fourel of the Institute of NeuroPhysiopathology, CNRS-Aix Marseille University, for their assistance with spinning-disk confocal microscopy.

