## Supplemental for "Oncogenic MAPK pathway activation disrupts Schwann cell fate commitment, inducing congenital and progressive neuropathy in mice"

#### Oncogenic MAPK pathway activation induces a congenital and progressively lethal rare neuropathy in mice

- Supplementary Methods
- Supplementary Figures and legends
- Supplementary references
- Legends to Supplementary Tables 1-3

### Supplementary Methods

#### Generation of transgenic *P0-Cre*; *Braf*$^{V600E/+}$ or *Krox20-Cre*; *Braf*$^{V600E/+}$ mice and genotyping

All mice were maintained on outbred backgrounds to maximize genetic heterogeneity by crosses over multiple generations to CD1/Swiss mice purchased from Janvier Laboratories (France). *Lox-Stop-Lox-Braf*$^{V600E/+}$ (*LSL-Braf*$^{V600E/+}$) mice, carrying a T1799A mutation in a floxed construct containing an alternative exon 15 within the *Braf* locus [1] (derived from RRID:IMSR_JAX:017928 but kindly provided by Prof. Richard Marais, University of Manchester, UK), were crossed to: (1) *P0(Mpz)-Cre* (derived from B6N.FVB-Tg(Mpz-cre)26Mes/J, RRID:IMSR_JAX:017927) [2,3] or (2) Krox20-Cre (RRID:IMSR_JAX:025744) [4] mice, to generate compound heterozygous *P0-Cre*$^{+/°}$; *Braf*$^{V600E/+}$ or *Krox20-Cre*$^{+/°}$; *Braf*$^{V600E/+}$ mice, called “mutant”; all other genotypes (WT$^{+/+}$, P0/Krox20-Cre$^{+/°}$ and *LSL-Braf*$^{V600E/+}$) were termed “control”.

Genotyping of *LSL-Braf*$^{V600E/+}$ x *Cre* descendants was performed using PCR primers to detect the *LSL-Braf*$^{V600E/+}$ construction allele: *LSL-Braf-Forward:* (5'-GCCCAGGCTCTTTATGAGAA-3') and *LSL-Braf-Reverse:* (5'-AGTCAATCATCCACAGAGACCT-3') as well as generic *Cre* primers for both driver lines: *Cre-Forward*: (5’-TGATGGACATGTTCAGGGATC-3’) and *Cre-Reverse*: (5’-CAGCCACCAGCTTGCATGA-3’). All PCR analyses were performed on DNA isolated from ear biopsies using Phire Tissue Master Mix PCR reagent (Thermo Fisher). The mice were kept in an animal facility with 12-hour light and 12-hour dark cycle and unrestricted access to other mice, water, and a conventional diet, with additional pellets provided in the bedding after weaning for cages with mutant animals. This animal study was reviewed and approved by the French accredited animal care and use committee (ACUC) C2EA-14 under the references 37-08102012 and 9522-2017040517496865v5.

#### Western Blot analyses

Mouse sciatic nerves were lysed in cold RIPA buffer (50 mM Tris (pH 8), 150 mM NaCl, 1% NP-40, 1% sodium deoxycholate, 0.1% sodium dodecyl sulfate (SDS) in the presence of 1X HALT protease inhibitor cocktail (1 mM AEBSF-HCl, 0.8 µM aprotinin, 50 µM bestatin, 15 µM E-64, 5 mM EDTA, 20 µM leupeptin, 10 µM pepstatin A; Thermo Fisher 78429). At 4°C, lysates were first sonicated for 3 cycles of 1 min pulse at 30 sec intervals in a Bioruptor (Diagenode) and then centrifuged at 4°C for 5 min at 1,000 *g*. Protein concentration was quantified by the DCProtein assay II (Bio-Rad 5000112) using an EnSpire Multimode plate reader (PerkinElmer) according to the manufacturer’s protocol. Lysates were diluted in NuPAGE LDS Sample Buffer (Invitrogen, Thermo Fisher), heated 5 min at 100°C under reducing and denaturing conditions in the presence of 25 mM dithiothreitol, and 20 µg of protein was separated on 12% SDS-PAGE gels. Proteins were transferred onto a 0.45 µm PVDF Immobilon-P membrane (Millipore, Merck) using the Pierce Power Blotter (Thermo Fisher). Membranes were blocked with 1X Tris-Buffered Saline-Tween (TBS-T; 50 mM Tris, 150 mM NaCl, 0.05% Tween-20, pH 7.6), containing 5% skim milk (1 hr at room temperature [RT]) and incubated with primary antibodies at 1:1000 dilution (overnight at 4°C): rabbit polyclonal anti-phospho (Thr202/Tyr204) Erk1/2 (#9101; Cell Signaling Technology, RRID:AB_331646), rabbit polyclonal anti-Erk1/2 (#9102; Cell Signaling Technology, RRID:AB_330744), rabbit monoclonal anti-Jun (clone 60A8) (#9165; Cell Signaling Technology, RRID:AB_2130165) or mouse monoclonal anti-GAPDH (#MAB374; Merck Millipore, RRID:AB_2107445). Alternately, the following primary antibodies were also used: mouse monoclonal anti-Shh (5E1, DSHB, RRID:AB_528466)[5], mouse monoclonal anti-Erbb3 (MAB348, R&D Systems, RRID:AB_358277), rabbit monoclonal anti-Egr2 clone EPR4004 (ab108399, Abcam, RRID:AB_10862073), mouse monoclonal anti-alpha-tubulin clone DM1A (ab7291, Abcam, RRID:AB_2241126), mouse monoclonal anti-vimentin (60330-1-Ig, Proteintech, RRID:AB_2881439 and mouse monoclonal anti-vinculin (V9264, Sigma-Aldrich, RRID:AB_10603627).

After washes, cells were incubated in the blocking solution with the HRP linked anti-mouse or anti-rabbit secondary antibody (respectively #7076 or #7074; Cell Signaling Technology, RRID:AB_330924 and RRID:AB_2099233) at 1:10,000 dilution (1 hr at RT in the dark). Blots were developed with the Super Signal West Pico or Femto PLUS Chemiluminescent Substrate (Thermo Fisher). Chemiluminescent signals were detected using a CCD camera and Syngene software (G:BOX). PageRuler Prestained protein ladder (#26616; ThermoFisher Scientific) was used as a molecular weight marker. Total protein band intensity was measured using NIH Image J software v.1.54f (https://imagej.net/ij/) and normalized to the corresponding level of Gapdh to control for variations in loading. The measurement of phosphorylated proteins was reported relative to the overall expression level of the protein of interest.

#### Culture and differentiation of human pluripotent stem cells (hiPSC)

This study was conducted in accordance with the Declaration of Helsinki [6], the Belmont report (Office for Human Research Protections, 2018), and the French law on bioethics 2021-1017, promulgated on August 2, 2021 (*LOI n° 2021-1017 du 2 août 2021 relative à la bioéthique*, 2021), consistent with our institutional review board protocols.

Two female patients with cardio-facio-cutaneous (CFC) syndrome bearing a heterozygous germline *BRAF* Q257R variant, or their legal guardians, provided written consent as approved by the UCSF Human Research Protection Program (CHR #10-02794), to produce the human induced pluripotent stem cell (hiPSC) lines, RMK0056C and RMK0138C, as described [9]. The AG08H and hFF15 clones of healthy donor male hiPSC lines were derived internally (CytoTune IPS 2.0 reprograming kit, Thermofisher) from commercial cell lines consented for research, AG08498 (RRID:CVCL_1Y51) and HFF-1 SCRC-1041 (RRID:CVCL_3285) respectively, as described [10]. After the latter had been assessed for absence of Sendai virus or mycoplasma and maintenance of pluripotency by our institutional Cell Reprogramming and Differentiation Facility (MaSC) in Marseille, France, all lines were initially grown in mTeSR1 medium (Stemcell Technologies) on human embryonic stem cell-qualified Matrigel™ (354277; BD Biosciences) before long-term storage over liquid nitrogen.

Established hiPSC lines were thereafter thawed and cultured on six-well plates coated with Synthemax II-SC Substrate (#734-2634; Corning) at a working concentration of 0.025 mg/mL and maintained in an undifferentiated state using StemMACS iPS-Brew XF medium (#130-104-368; MACS Miltenyi Biotec). The hiPSC lines were incubated at 37°C and 5% $CO_2$ and media changed daily. When cells reached 60-80% confluence, they were passed using enzyme-free ReleSR (#05872; StemCell Technologies), adding 10 µM of ROCK inhibitor Y-27632 (#SCM75; Sigma-Aldrich) for the first 24 hours after passage.

For the directed differentiation of hiPSC into SC, we followed a published protocol [11]. Differentiation of hiPSCs into SC was initiated when both control and mutant *BRAF* Q257R hiPSCs reached a confluence of 60%. Subsequently, cells were passaged every 6 days until day 31, at which point cells were passaged upon reaching 90% confluence, not necessarily synchronized between the cell lines. During the differentiation period, cell passaging was conducted as follows: at a 1:10 split on the 6th day, at a 1:6 split on the 12th and 18th days, and at a 1:4 split on the 24th day. From day 31 onward, cell passaging was carried out at a 1:2 split to maintain an adequate cell density and prevent cellular death.

#### Total RNA isolation

Sciatic nerves from WT or *Braf*$^{V600E/+}$ animals of both sexes, and pellets of WT or *BRAF*$^{Q257R/+}$ hiPSC-derived SCP and immature Schwann cells (iSC) were stored at -80°C. Sciatic nerves on postnatal days 5 (P5) or 21 (P21) were lysed using ceramic beads in the FastPrep-24 homogenizer (MP Biomedicals) at 6 m/s for 50 sec and total RNA extracted with the RNeasy Lipid Tissue Mini Kit (#74804; Qiagen). For SCP (D18) and iSC (D31) differentiated from hiPSCs, total RNA was extracted using the NucleoSpin RNA Plus S kit (#740984; Macherey-Nagel). RNA concentrations were measured on a NanoDrop (NanoDrop TM 1000 Spectrophotometer; Thermo Fisher Scientific).

#### Library preparation and RNA-sequencing

RNA-sequencing and bioinformatics analysis was carried out at the GBiM Genomics and Bioinformatics facility from the U1251/Marseille Medical Genetics laboratory. We undertook three different RNAseq analyses. The first was carried out on entire pituitary glands microdissected from four replicate P5 (three female and one male control, four female Mpz-*Braf*$^{V600E+/-}$ mutant) and two replicate P21 (two female control, one female and one male mutant) glands for 12 total samples (Supplementary Table S1). The second was performed with total RNA samples obtained from microdissected thigh segments of P5 or P21 sciatic nerves collected from either control or mutant mice (16 samples in total, Supplementary Table S2). At P5, these comprised three female and one male control nerves for both genotypes. At P21, there were one female and three male controls *vs* one male and three female mutant nerves. The third RNAseq was performed on two or three replicates from hiPSC-derived SCP differentiated from two WT (hFF15 and AG08H, derived in-house) or two *BRAF*$^{Q257R/+}$ mutant female (RMK0056C and RMK0138C; [9] hiPSCs (29 samples in total, Supplementary Table S3). Following RNA extraction, we evaluated quality using a Bioanalyzer (Agilent Technologies). Total RNAs with RNA Integrity Numbers (RIN) exceeding 8 were used. For each sample, a library was prepared from 1 µg of total RNA for whole sciatic nerves and pituitaries or from 300 ng of total RNA for differentiated cells, after capture of polyadenylated RNA using the KAPA mRNA HyperPrep kit (#KR1352-v7.21; Roche). The individual libraries were assessed using a Qubit™ (ThermoFisher) and the Agilent Bioanalyzer dsDNA High Sensitivity Kit (Agilent Technologies). We sequenced at 2*76 basepairs (mouse sciatic nerves, P5 pituitaries) or at 2*101 base-pairs (from hiPSC-derived SC, P21 pituitaries) on a NovaSeq 6000 (Illumina).

#### Data processing and differential gene expression analysis (DGE)

The quality of sequencing reads was assessed using fastQC (v0.11.5) (https://www.bioinformatics.babraham.ac.uk/projects/fastqc/). Raw sequencing reads were aligned on the GRCm39 mouse reference genome or on the GRCh38 human reference genome using STAR (v2.7.2b). Bam files were indexed and sorted using Samtools (v1.7). After mapping, the number of reads per feature (Ensembl or GENCODE v34 annotations for mouse or human data, respectively) was determined using StringTie v2.1.6.

Genes differentially expressed between conditions were identified using a Wald test in the DESeq2 package (v1.34.0). Differentially expressed transcripts with an adjusted P-value for FDR (False Discovery Rate) below 0.05 were considered significant (see Statistics, below). Relative expressions of the most variable features between samples were plotted as heatmaps using the Pheatmap R package. Differentially Expressed Genes (DEGs) were visualized in the form of volcano plots, representing the Log2FC (log2 of the expression -fold change) and the adjusted P-value, prepared using the EnhancedVolcano R package [12].

#### Nerve and fiber immunostaining

For immunofluorescence analyses, sciatic nerves were isolated from mice, fixed for 30 min (2 hr for mutants) in 4% paraformaldehyde in PBS at pH 7.4 and washed in PBS. Teased fibers were separated with insect pins onto Superfrost Plus glass slides, or whole nerves were embedded in Paraplast Plus for classical histological sections at 6 µm. Slides were then dried overnight at RT and stored at 4°C for wax sections or at -20°C. Teased sciatic nerves were permeabilized in acetone for 10 min at -20°C, washed in PBS, non-specific binding blocked for 1 hr at room temperature in PBS containing 5% w/v fish skin gelatin and 0.5% Triton-X100, then incubated with primary antibodies diluted in the blocking solution overnight at 4°C. Wax sections were deparaffinated in xylene and rehydrated, treated with freshly prepared 50 mM glycine for 20 minutes, blocked in PBS with 0.5% Tween-20 and 2% donkey serum, and incubated overnight with primary antibodies diluted in the blocking buffer, without heat-inactivated antigen retrieval. After multiple PBS/0.5% Tween-20 rinses, slides were incubated with the appropriate Alexa Fluor-coupled secondary antibody (Invitrogen/ThermoFisher) and DAPI (4',6-diamidino-2-phenylindole) diluted respectively to 1/500 and 300 nM in the blocking solution for 1 hr at room temperature. Teased sciatic nerves were washed and mounted in Prolong Diamond medium (#P36961; Invitrogen) while sections were mounted in Fluoromount-G™ (ThermoFisher). Image stacks of teased fibers were acquired using a super-resolution spinning disk confocal microscope (Nikon SoRa) and NIS-Elements software; a Zeiss LSM800 with Airyscan and Zen software v2.3 acquired confocal optical slices of nerve sections.

The following antibodies were used for immunolabeling tissues: rat monoclonal to MBP at 1/250 (MAB386, Millipore, RRID:AB_94975); rat monoclonal to Ki-67 at 1/100 (SolA15, ThermoFisher, RRID:AB_10854564); rabbit polyclonal to Caspr at 1/50 (ab34151, Abcam, RRID:AB_869934), rabbit monoclonal to Sox10 at 1/250 (AC-0237, Cell Marque, RRID:AB_2941085); rabbit polyclonal to Pmp2 at 1/100 (12717-1-AP, Proteintech, RRID:AB_2166978); mouse monoclonal to Erbb3 at 1/100 (MAB348, R&D Systems, RRID:AB_358277); rabbit polyclonal to S100b at 1/1000 (HPA015768, Atlas Antibodies, RRID:AB_1856538); and mouse monoclonal to Shh at 1/100 (5E1, DSHB, RRID:AB_528466)[5]. Alexa Fluor 488-coupled phalloidin was also used at a 1/500 dilution to label filamentous (F)-actin (A12379, ThermoFisher).

#### Transmission electron microscopy (TEM) and image analysis

Mouse sciatic nerve samples were dissected and fixed in fresh ice-cold 2% PFA + 2.5% glutaraldehyde in phosphate buffered saline at pH 7.4 for 7 hrs. After three washes in 0.1M sodium cacodylate buffer, nerves were post-fixed with 1% osmium tetroxide ($OsO_4$) for 2 hr on ice. The nerves were rinsed in distilled water and contrasted in aqueous 1% uranyl acetate solution overnight at 4°C. Samples were then dehydrated in a series of ethanol solutions of increasing concentrations for 20 min each and infiltrated with Epon resin in acetone (in ratios of 1:3, 2:2 and 3:1) for 2 hr each. Finally, nerves were embedded in fresh resin overnight and polymerized at 60°C for 48 hr. Semi-thin sections at 1 µm were stained with toluidine blue. Ultra-thin sections for TEM were mounted on copper grids coated with Formvar membrane and contrasted with 2% uranyl acetate for 10 min and lead citrate for 5 min before imaging on an FEI Morgagni 120 kv electron microscope.

#### Analysis of sciatic nerve morphology

MyelTracer software [13] was used on 8-bit TEM images to determine the *g*-ratio (axon diameter/fiber diameter), axon diameter, and the distribution of axons according to the thickness of the myelin sheath in control vs mutant sciatic nerves at P5 (n = 3 individuals per group) and at P28 (n = 2 individuals per group). The number of axons was counted using ImageJ software on lower magnification TEM images of the sciatic nerve, comprising n = 3 and 3, n=5 and 4, and n=2 and 2 controls *vs* mutants at P5, P21 and P28, respectively. Analyses were conducted with average equal sex ratios.

#### Enrichment analysis

Expression-dependent enrichment analysis was conducted using Gene Set Enrichment Analysis (GSEA) software v4.3.2 [14] with the mouse Wikipathway database to compute pathway enrichment scores in P0(Mpz)-$\textit{Braf}^{V600E/+}$ (Mpz-*Braf*) sciatic nerves compared to controls. The same analysis was carried out using the human Reactome database for $\textit{BRAF}^{Q257R/+}$ hiPSC-derived SCP or iSC in comparison to control-differentiated hiPSCs.

Expression-independent enrichment analysis was performed using g:Profiler [15] or Enrichr-KG [16], interrogating the Wikipathway, KEGG, Reactome and Gene Ontology datasets.

### Statistics

Statistical analyses were conducted using Prism v10.2.0 for Windows (GraphPad Software, Boston, Massachusetts USA, www.graphpad.com). Two-sided comparisons were assessed using unpaired Mann-Whitney or two-way ANOVA tests for non-parametric tests. For multiple comparisons, two-way ANOVA tests followed by Šidák's post-hoc method were used, or Student t-tests followed by Benjamini-Hochberg adjustment to control the False

Discovery Rate (FDR) in DESeq2. Mendelian segregation was assessed with a chi-square contingency test. Data are presented as the mean ± standard error of the mean (s.e.m). $p > 0.05$ was considered as non-significant (ns), (*) $p < 0.05$; (**) $p < 0.01$; (***) $p < 0.001$; (****) $p < 0.0001$.

#### Supplementary Figures and Legends

### Supplementary Figure S1

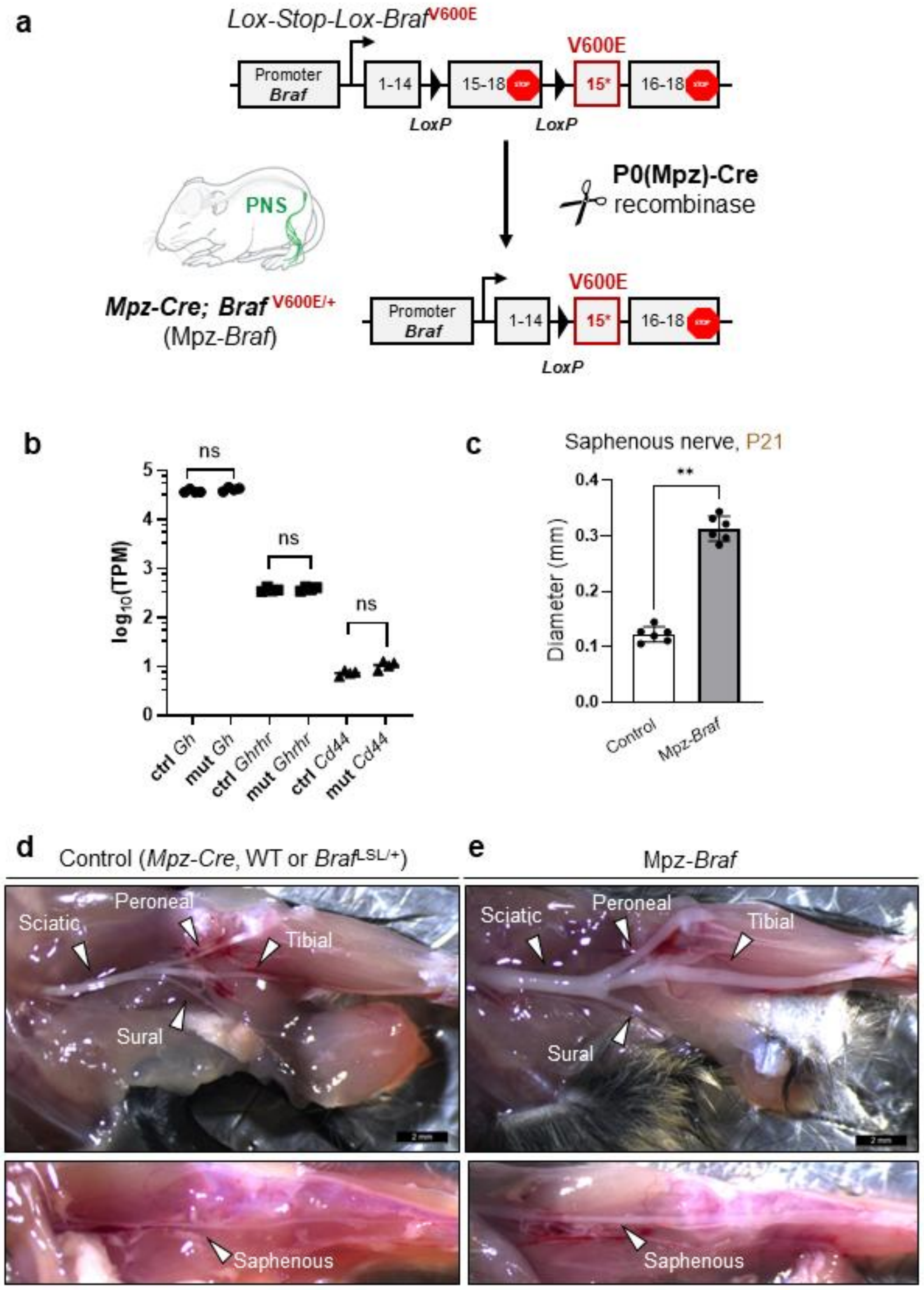

#### Supplementary Figure S1.

Genotype and phenotype of mouse crosses. **a**. to produce Mpz-Cre; Braf(V600E/+) offspring, Mpz-Cre (also known as P0-Cre) driver males were crossed to Braf(LSL-V600E/+) mice, sometimes also carrying a floxed tdTomato allele for simultaneous lineage tracing of recombined cells. **b**. At P5, bulk RNA-sequencing showed no significant differences after p-value adjustment for multiple hypothesis testing in transcription of *Gh*, *Ghrhr* or *Cd44* genes, except for a predicted X-linked pseudogene, *Gm14584* ($p<0.03$). For these and other transcripts at both P5 and P21, see Table S1. **c**. The saphenous sensory nerve at P21, like the trigeminal ganglion and mixed sensory-motor nerves, was significantly wider than its littermate counterpart ($p<0.01$). The widespread nature of nerve enlargement accompanied by muscle wasting is visible in these exemplary dissections to expose sensory or mixed sensory-motor nerves in control (**d**) versus mutant (**e**) mouse legs at P21.

### Supplementary Figure S2

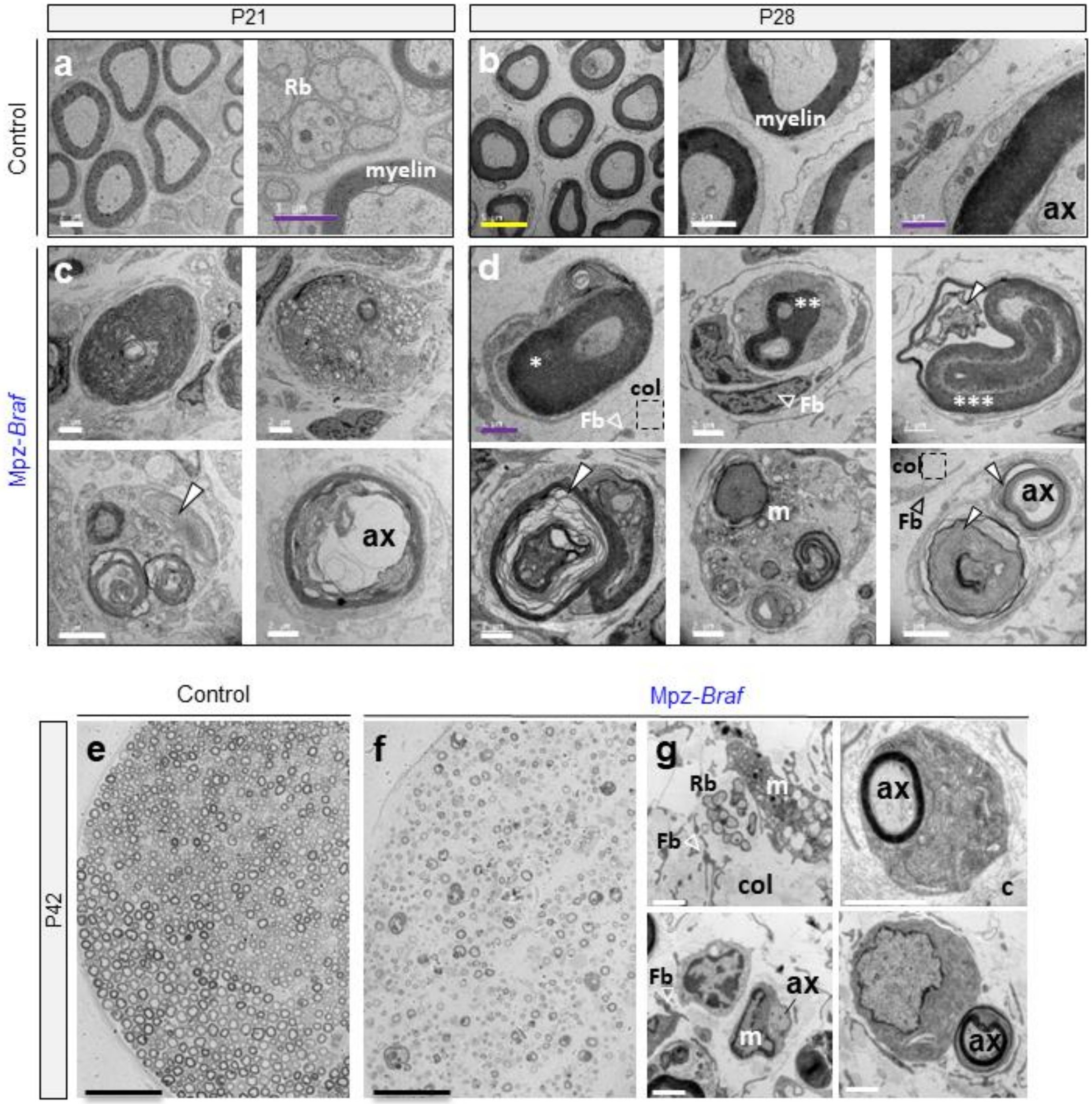

#### Supplementary Figure S2.

**a-d** and **g.** Transmission electron micrographs (TEM) of cross-sections of control or Mpz-Braf sciatic nerves on postnatal days (P)21, P28 and P42. Control nerves show myelinated fibers of varying diameters (**a, b**) and non-myelinated small axons organized in Remak bundles (Rb, **a, g**). In contrast, myelin was observed at various stages of degradation at all stages post-weaning (white filled arrowheads), with the exception of hypermyelination of small-caliber axons surrounded by “onion bulb” formations (asterisks). Semi-thin resin sections stained with toluidine blue (**e, f**) at P42 showed a dramatic overall reduction of mutant myelinated axons (ax) and (**g**) in TEM, abnormal myelin structures but maintenance of some Remak bundles, accompanied by abundant extracellular collagen (col) from intraneurial fibroblasts (Fb) as well as macrophages (m) containing myelin debris. Scale bars: (**a-d**) violet = 1 μm, white = 2 μm, yellow = 5 μm; **e, f** = 50 μm and **g** = 2 μm.

### Supplementary Figure S3

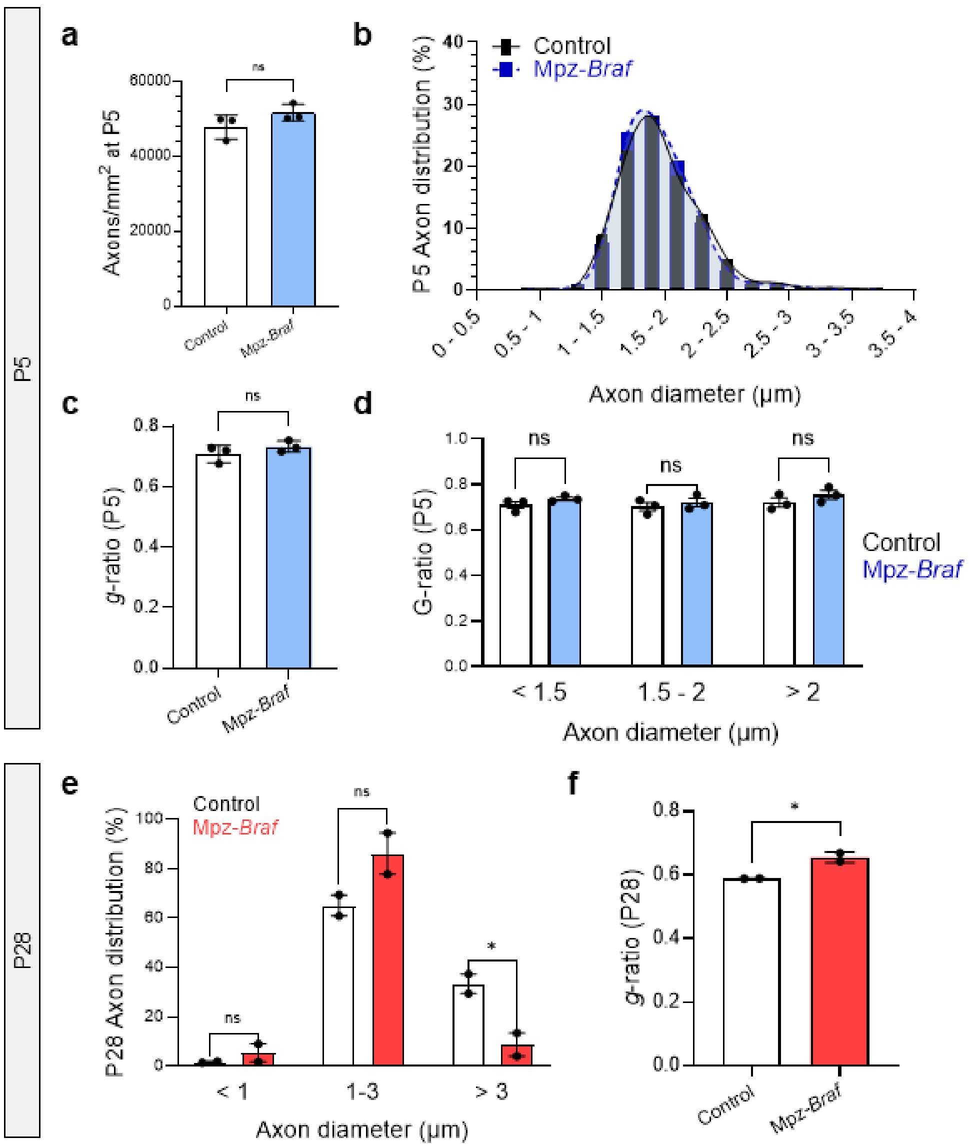

#### Supplementary Figure S3.

**a**) No significant difference in axon density in P5 mutant mice from littermate controls (n=3 each) in cross-section of the sciatic nerve. **b**) Distribution of axon diameters at P5 are also not significantly different between Mpz-*Braf* and control littermate mice. **c**) g-ratios of the myelin sheaths around the axons in the cross-sections analyzed in (**a**) are not significantly different when analyzed together. (**d**) Same data, binned by size to show no distinction by axon caliber. At P28 (e) the data binned by caliber shows, in contrast, that large-caliber axons are preferentially less frequent. (**f**) On average, g-ratios at P28 are greater, indicating thinner myelin sheaths overall.

### Supplementary Figure S4

**a** Control P42

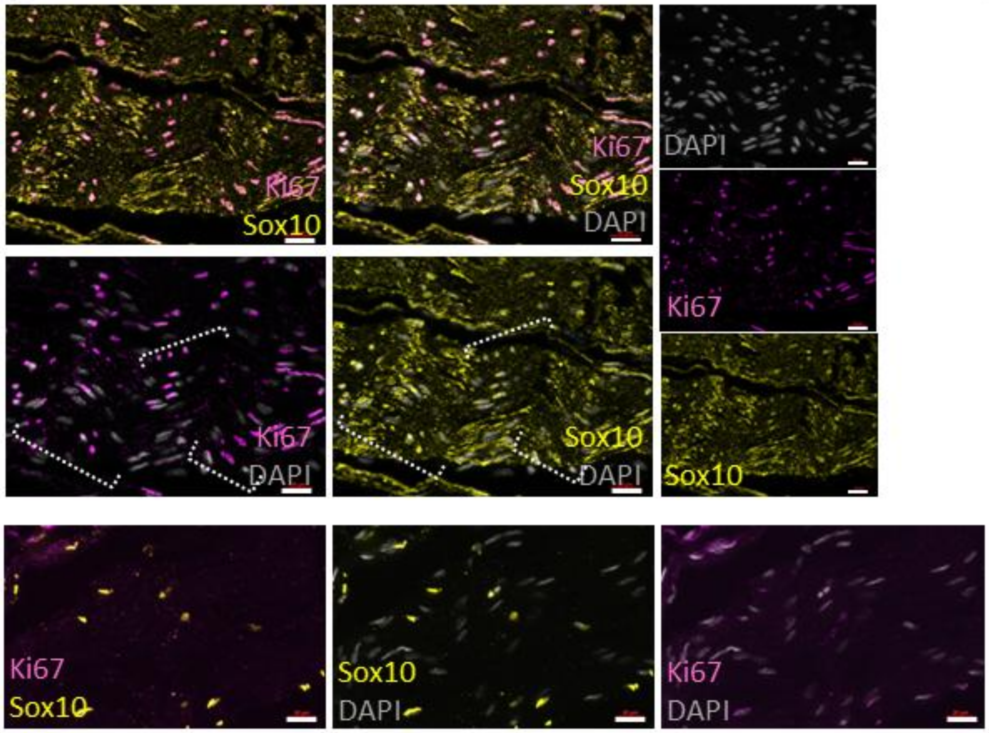


**b** Mpz-*Braf* P42

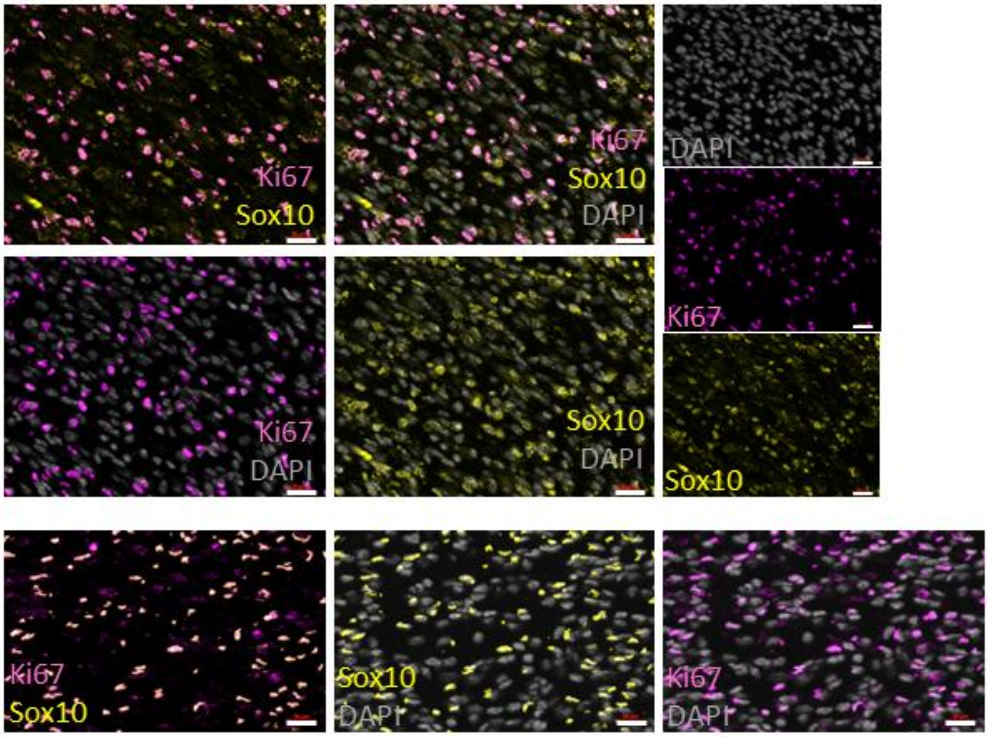

#### Supplementary Figure S4.

Co-immunostaining of Sox10, a SC-specific transcription factor in peripheral nerves, and Ki67, a marker of non-quiescent cells having bypassed the G0/G1 cycle checkpoint (Miller et al., 2018). **a**. Sections of p42 sciatic nerves from two control mice from distinct litters (upper and lower frames), showing Fontana band organization (brackets) with SC nuclei alignedalong nerve fibers. Many but not all nuclei (DAPI, gray) express Sox10 (yellow), but these all appear to co-express at least some Ki67 (magenta). **b**. Sections of mutant P42 sciatic nerves from two Mpz-*Braf* mice from distinct litters. Nuclei are present at a higher density than in controls under analogous processing conditions. All Sox10+ nuclei continue to co-express Ki67 as do some Sox10-negative cells, potentially corresponding to fibroblasts or tissue-resident macrophages. Slides were processed simultaneously with the same batches of primary and secondary antibody dilutions in a single experiment, along with no-primary antibody controls, and photographed under identical exposure and adjustment parameters with Zeiss Zen software on a confocal microscope to ensure comparability.

### Supplementary Figure S5

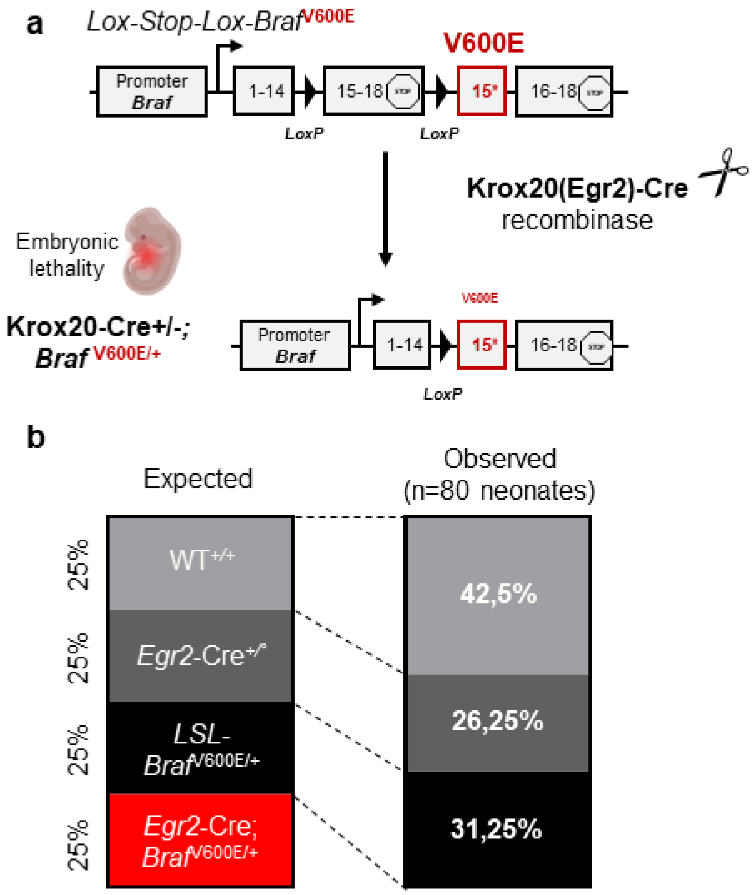


| | *Egr2-Cre*$^{+/°}$ x *LSL- Braf*$^{V600E/+}$ | | | | |
|---|---|---|---|---|---|
| | WT$^{+/+}$ | *Egr2-Cre*$^{+/°}$ | *LSL-Braf*$^{V600E/+}$ | *Egr2-Cre; Braf*$^{V600E/+}$ | Total |
| Expected | 20 (25%) | 20 (25%) | 20 (25%) | 20 (25%) | 80 (100%) |
| Observed | 34 (42,5%) | 21 (26,25%) | 25 (31,25%) | **0 (0%)** | 80 (100%) |

#### Supplementary Figure S5.

**a**) Schematic representing crossing the lox-stop-lox(LSL)-*Braf*$^{V600E}$ mouse line to the Egr2(Krox20)-*Cre* driver line. **b**) The distribution of genotypes at birth resulting from the mating of heterozygous LSL-*Braf*$^{V600E/+}$ mice with Egr2-*Cre* mice was significantly different from expected ($p < 0.0001$, chi-square 24.21, df 3), demonstrating fully penetrant prenatal lethality (red rectangles).

### Supplementary Figure S6

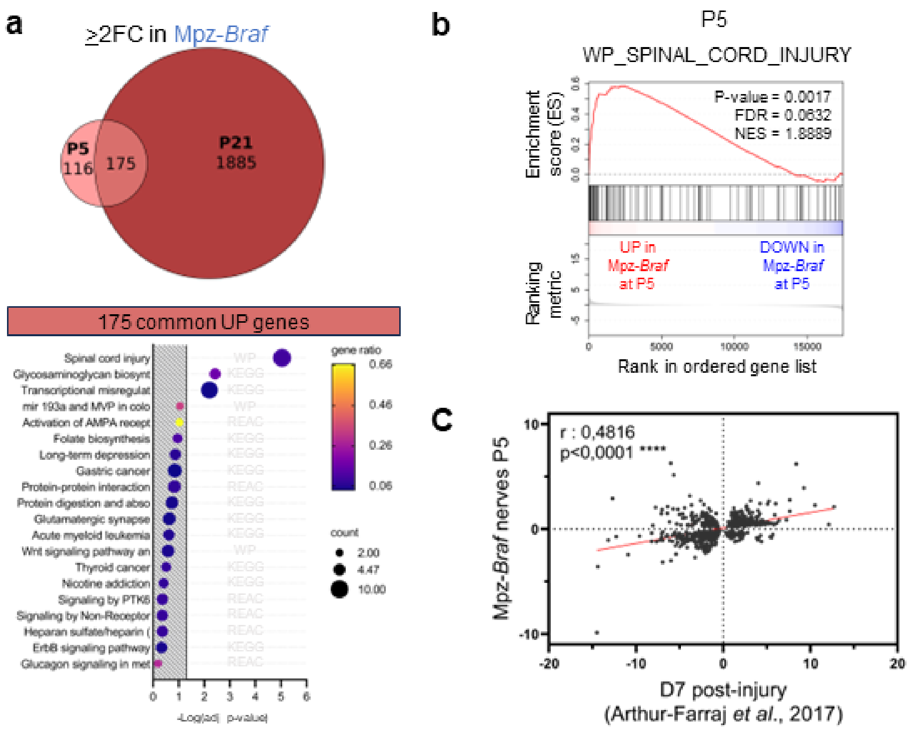


#### Supplementary Figure S6.

**a**) Euler diagram of commonly or stage-specific significantly upregulated (greater than two-fold, padj < 0.05) transcripts in RNAseq from Mpz-*Braf* mutant mouse sciatic nerves, relative to stage-matched controls. 175 transcripts were commonly upregulated at both stages, and were annotated by terms from Wikipathways (WP), KEGG or Reactome (REAC) databases with the indicated significance, where terms beyond the hatched areas were significantly enriched at an adjusted p<0.05. **b**) GSEA analysis also showed that even at P5, a significant number of all enriched transcripts were annotated in the WP2431 Wikipathway “Spinal cord injury”. **c**). A highly significant direct correlation was found of P5 Mpz-*Braf* DEGs in common with DEGs in a mouse sciatic nerve injury model after a week post-injury, during the regeneration phase (cf. Arthur-Farraj *et al.* 2017).

### Supplementary Figure S7

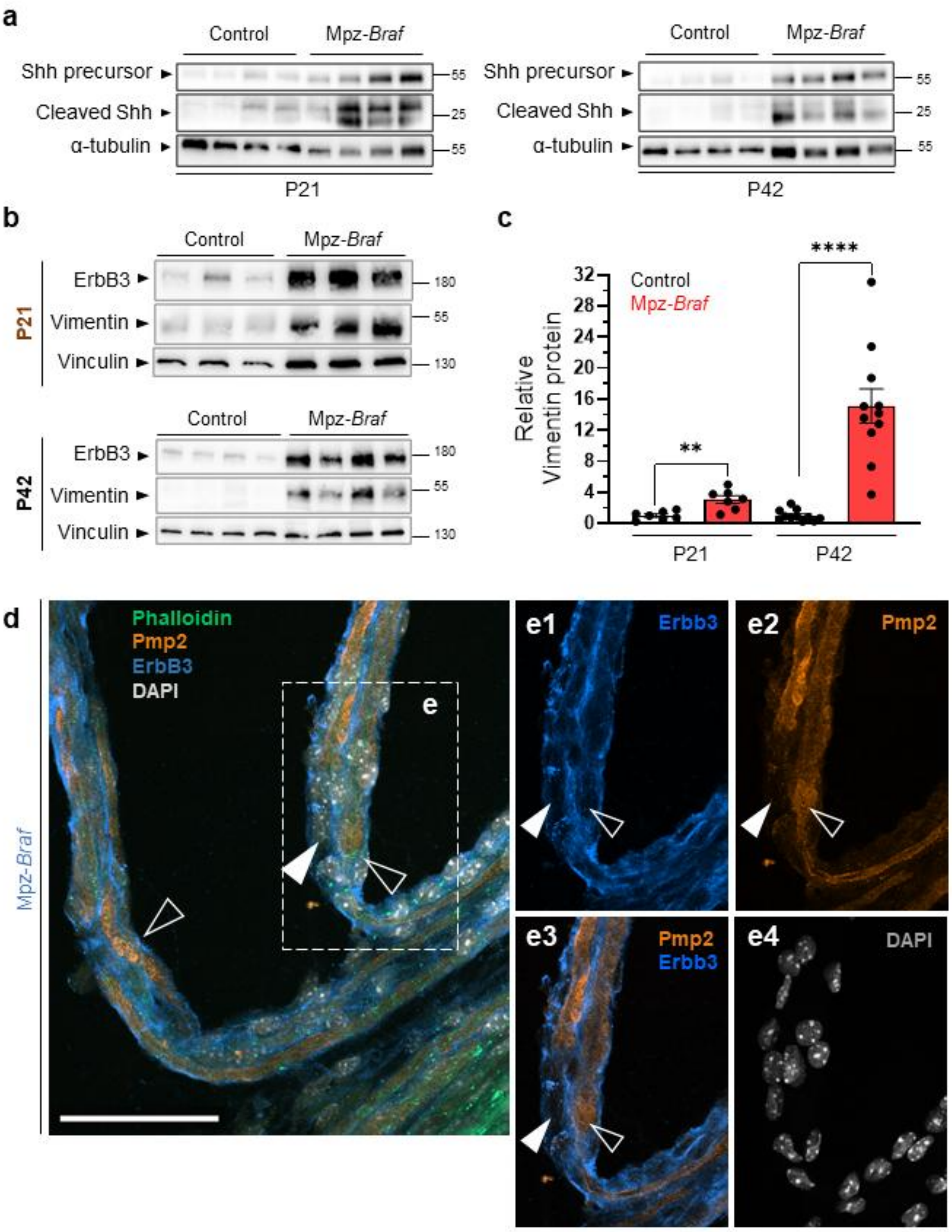

#### Supplementary Figure S7.

**Supplementary Figure 7. a**) Among the top transcripts upregulated at P5, Shh as assessed by Western blot of Mpz-*Braf* and control sciatic nerves from four mice each at P21 was significantly more expressed in both total and active cleaved forms in mutant nerves, and even more so by P42. α-tubulin was used as a loading control (see **c**). **b**) Western blots of sciatic nerve extracts from three mice at P21 and three at P42 to assess expression of ErbB3, vimentin and vinculin as a high molecular weight loading control. **c**) Vimentin was not an appropriate loading control for mid-range molecular weights, as when it was normalized to vinculin, it became increasingly translated in mutant sciatic nerves at P21 and P42. This observation is associated with increased ECM deposition in the same nerves (*cf*. Fig. 2). **d**) Immunodetection of the myelin protein Pmp2 and the Nrg1 receptor ErbB3 in teased sciatic nerve fibers from a Mpz-Braf mouse at P21, with green F-actin staining of axons with phalloidin and white nuclear staining with DAPI. **e**) Single (**e1, e2, e4**) or two combined channels (**e3**) to show separate components in the fibers boxed in (**d**), where some Schwann cells continue to express both proteins, albeit unevenly distributed (black arrows) and others show diminished Erbb3 and undetectable Pmp2 (white arrows), indicating demyelination.

#### Supplementary Table 1.

RNAseq of P5 and P21 pituitaries from Mpz-*Braf* and control littermate mice.

#### Supplementary Table 2.

RNAseq of P5 and P21 sciatic nerves from Mpz-*Braf* and control littermate mice.

#### Supplementary Table 3.

RNAseq of hiPSC undergoing a directed differentiation protocol towards a Schwann cell phenotype on days (D)18 and 31 from two CFC patient cell lines, abbreviated 56C and 138C. Yellow tabs analyze both patients relative to both control lines at the same time points, while intermediate tabs show DEGs of each cell line relative to both control lines at the same time points. In red and blue are Venn diagrams to examine which DEGs are preferentially up- or downregulated, respectively, by one or the other individual lines or timepoints or at the intersection of all mutant versus control lines at both timepoints (D18_56C_UP ∩ D18_138C_UP ∩ D31_56C_UP ∩ D31_138C_UP and D18_56C_DOWN ∩ D18_138C_DOWN ∩ D31_56C_DOWN ∩ D31_138C_DOWN). For example, *EGR2* is commonly downregulated in both CFC cell lines and at both d18 and d31, relative to control cells.

### Supplementary References

RASopathies are heterogeneous developmental syndromes that preferentially affect epidermis and pigmentation; cardiovascular, craniofacial and skeletal systems; and neurological or endocrine functions impacting body size and fertility [17–20].

Activating variants may be expressed constitutionally or as mosaic lineage-specific (somatic) RASopathies [21]. Most constitutional and somatic RASopathies associate with specific cancer predispositions, particularly when syndromic [22,23].

Oncogenic variants in *BRAF*, particularly encoding changes to valine (p.V)600, drive aggressive forms of melanoma as well as many other cancers [24,25].

BRAF p.V600E yields an active, monomeric kinase, decoupled from RAS-mediated transduction [26,27]. In contrast, less-activating *BRAF* variants such as p.Q257R underlie a growing list of congenital RASopathies [22,28]. These include the germline disorders cardio-facio-cutaneous syndrome (CFC; ORPHA:1340) and Noonan syndrome with (ORPHA:500) or without (ORPHA:648) multiple lentigines. Their common clinical features include reduced postnatal growth, dysmorphic facial features, melanocytic nevi or other pigmented lesions, congenital heart defects, and tumor predispositions [29–31].

Recent reports describe CFC and Noonan syndrome patients with early-onset neuropathies characterized by pain, sensory loss and/or muscle wasting [32–34], resembling severe, early-onset Charcot-Marie-Tooth (CMT) syndromes [35,36]. Peripheral nervous system (PNS) function relies entirely on Schwann cells (SC), which provide physical and trophic nerve support while insulating large, fast-conduction axons [37–39]. The PNS — including SC, perineurial stroma and sensory/autonomic neurons — derives from multipotent embryonic neural crest (NC) cells, which also generate craniofacial and cardiovascular connective tissues, neuroendocrine cells and melanocytes [39–42]. Mek1, the primary MAP2K activating Erk1/2, is essential for nerve myelination during maturation, but its sustained overactivation in adult SC induces murine neuropathy [43–46].

*Mpz-Cre*$^{+/\circ}$ mice, which transcribe *Mpz* (encoding myelin protein zero, or P0) as early as embryonic day 12 in neural crest cells [2,47].

Gh binds to peripheral Ghr, a type I cytokine receptor, to regulate production of its effector, insulin-like growth factor 1 (Igf1) in the liver and other sites, exerting thereby anabolic effects on bone and muscle [48].

Strikingly broad nerves were visible at autopsy in all Mpz-*Braf* mice at P5 or P21, lacking the structural bands of Fontana [49].

Ki67 is a cell cycle progression marker [50,51].

Strongly downregulated DEGs at P21 included *Hmgcr* (11.5-fold reduction), encoding the rate-limiting enzyme in cholesterol biosynthesis [52], and *Fa2h* (7.9-fold reduction), encoding fatty acid 2-hydroxylase, a key enzyme in myelin glycosphingolipid synthesis [53].

*Ptpn11* is a Noonan syndrome RASopathy-associated gene [54].

This fully penetrant embryonic lethality, likely due to broader expression of *Egr2* beyond SCs [55,56], prevented further postnatal myelination comparisons (**Fig S5B**).

To determine whether Jun-dependent repair ligand effectors were also upregulated, we examined *Shh* (Sonic hedgehog) and the neurotrophic factor *Gdnf* (glial cell line-derived neurotrophic factor) [57].

Cleaved Shh is critical for multipotent NC maintenance [58].

Nrg1-III and the Gdnf family member Artemin (Artn), secreted by Jun-positive repair SCs to promote axon outgrowth [59,60], were upregulated at P5 (*Nrg1*) and P21 (*Nrg1*, *Artn*) (**Fig. 7c**). The Nrg1 receptor *ErbB3*, a nerve damage hallmark [61], was transcriptionally and translationally upregulated over time.

Vimentin (*Vim*), an intermediate filament upregulated in injured axons [62], also increased over time (**Fig. S7c; Table S2**). Despite repair program activation, progressive axonal degeneration in mutants implies that sustained MAPK signaling is counterproductive for SC-mediated trophic activity [63] *in vivo*.

To study a RASopathy-associated variant [64], we differentiated human induced pluripotent stem cells (hiPSCs) from two unrelated CFC patients.

Reactome [65] pathway analysis.

*LAMA2*, encoding the alpha-2 chain of laminin-211, is a component of SC basal lamina necessary to restrain small-caliber fiber hypermyelination [66].

*POU3F2* is a TF transiently required in neonatal murine SC [67].

Prolonged postnatal MAPK signaling in immature SC, both in mice and humans, sustains transcriptional programs typical of other neural crest lineages [42,68].

A key phenotype was rapid onset of hyperplastic neuropathy, reminiscent of "hypertrophic neuropathy" described in case reports [32,34,69–72]. This phenotype had not been reported in prior systemic RASopathy animal models [73].

Like other RASopathy models (e.g. germline mutations in *Kras* [74], *Braf* [75–78], *Raf1* [79], *Ptpn11* [80] or *Sos1* [81]), the Mpz-*Braf* mouse presents early postnatal failure to thrive. Severe feeding difficulties are common in *BRAF*-associated germline RASopathies [32,82–84].

Postnatal *Igf1*-null mice, which survive at low rates, exhibit severe growth retardation [85,86], while liver-specific Igf1 rescue restores both body size and fertility [87]. The fully penetrant cryptorchidism in male Mpz-*Braf* but not *Igf1*-null mice, despite no *Mpz* expression in germ cells, suggests a neurogenic mechanism. Cryptorchidism is common in RASopathies [19,88], and genitofemoral nerve damage in rats causes testicular ascent [89]. This supports a role for peripheral nerve myelination in maintaining testicular descent and fertility [90,91].

--


1. Mercer, K. *et al.* Expression of endogenous oncogenic V600EB-raf induces proliferation and developmental defects in mice and transformation of primary fibroblasts. *Cancer Res.* **65**, 11493–11500 (2005).
2. Feltri, M. L. *et al.* A novel P 0 glycoprotein transgene activates expression of lac Z in myelin-forming Schwann cells. *Eur. J. Neurosci.* **11**, 1577–1586 (1999).
3. Feltri, M. L. *et al.* P0-Cre transgenic mice for inactivation of adhesion molecules in Schwann cells. *Ann. N. Y. Acad. Sci.* **883**, 116–123 (1999).
4. Voiculescu, O., Charnay, P. & Schneider-Maunoury, S. Expression pattern of a Krox-20/Cre knock-in allele in the developing hindbrain, bones, and peripheral nervous system. *Genes. N. Y. N 2000* **26**, 123–126 (2000).
5. Ericson, J., Morton, S., Kawakami, A., Roelink, H. & Jessell, T. M. Two critical periods of Sonic Hedgehog signaling required for the specification of motor neuron identity. *Cell* **87**, 661–673 (1996).
6. The World Medical Association, (WMA). WMA Declaration of Helsinki. *Ethical Principles for Medical Research Involving Human Subjects* https://www.wma.net/policies-post/wma-declaration-of-helsinki-ethical-principles-for-medical-research-involving-human-subjects/ (2022).
7. Office for Human Research Protections, (OHRP). The Belmont Report. *Ethical principles and guidelines for the protection of human subjects of research* https://www.hhs.gov/ohrp/regulations-and-policy/belmont-report/read-the-belmont-report/index.html (2018).
8. Journal Officiel, F. *LOI n° 2021-1017 du 2 août 2021 relative à la bioéthique*. (2021).
9. Yeh, E. *et al.* Patient-derived iPSCs show premature neural differentiation and neuron type-specific phenotypes relevant to neurodevelopment. *Mol. Psychiatry* **23**, 1687–1698 (2018).
10. Dion, C. *et al.* SMCHD1 is involved in de novo methylation of the DUX4-encoding D4Z4 macrosatellite. *Nucleic Acids Res.* **47**, 2822–2839 (2019).
11. Hörner, S. J. *et al.* hiPSC-Derived Schwann Cells Influence Myogenic Differentiation in Neuromuscular Cocultures. *Cells* **10**, 3292 (2021).
12. Blighe, K., Rana, S. & Lewis, M. Blighe K, Rana S, Lewis M (2023). EnhancedVolcano: Publication-ready volcano plots with enhanced coloring and labeling. R package version 1.20.0. (2023).
13. Kaiser, T. *et al.* MyelTracer: A Semi-Automated Software for Myelin g-Ratio Quantification. *eNeuro* **8**, ENEURO.0558-20.2021 (2021).
14. Subramanian, A. *et al.* Gene set enrichment analysis: A knowledge-based approach for interpreting genome-wide expression profiles. *Proc. Natl. Acad. Sci.* **102**, 15545–15550 (2005).
15. Kolberg, L. *et al.* g:Profiler—interoperable web service for functional enrichment analysis and gene identifier mapping (2023 update). *Nucleic Acids Res.* **51**, W207–W212 (2023).
16. Evangelista, J. E. *et al.* Enrichr-KG: bridging enrichment analysis across multiple libraries. *Nucleic Acids Res.* **51**, W168–W179 (2023).
17. Gualtieri, A. *et al.* Activating mutations in BRAF disrupt the hypothalamo-pituitary axis leading to hypopituitarism in mice and humans. *Nat. Commun.* **12**, 2028 (2021).
